# Relative contributions of sex hormones, sex chromosomes, and gonads to sex differences in tissue gene regulation

**DOI:** 10.1101/2021.07.18.451543

**Authors:** Montgomery Blencowe, Xuqi Chen, Yutian Zhao, Yuichiro Itoh, Caden McQuillen, Yanjie Han, Benjamin Shou, Rebecca McClusky, Karen Reue, Arthur P. Arnold, Xia Yang

## Abstract

Sex differences in physiology and disease in mammals result from the effects of three classes of factors that are inherently unequal in males and females: reversible (activational) effects of gonadal hormones, permanent (organizational) effects of gonadal hormones, and cell-autonomous effects of sex chromosomes, as well as genes driven by these classes of factors. Often, these factors act together to cause sex differences in specific phenotypes, but the relative contribution of each and the interactions among them remain unclear. Here, we used the Four Core Genotypes (FCG) mouse model with or without hormone replacement to distinguish the effects of each class of sex-biasing factors on transcriptome regulation in liver and adipose tissues. We found that the activational hormone levels have the strongest influence on gene expression, followed by the organizational gonadal sex effect and, lastly, sex chromosomal effect, along with interactions among the three factors. Tissue specificity was prominent, with a major impact of estradiol on adipose tissue gene regulation, and of testosterone on the liver transcriptome. The networks affected by the three sex-biasing factors include development, immunity and metabolism, and tissue-specific regulators were identified for these networks. Furthermore, the genes affected by individual sex-biasing factors and interactions among factors are associated with human disease traits such as coronary artery disease, diabetes, and inflammatory bowel disease. Our study offers a tissue-specific account of the individual and interactive contributions of major sex-biasing factors to gene regulation that have broad impact on systemic metabolic, endocrine, and immune functions.

## INTRODUCTION

Females and males differ in the incidence, progression, and genetic risks of many diseases, such as metabolic disorders including obesity, non-alcoholic fatty liver disease, and diabetes (Arnold 2010; Clayton and Collins 2014; Rask-Andersen et al. 2019). Thus, one sex may have endogenous protective or risk factors that could become targets for therapeutic interventions. Current sexual differentiation theory suggests that three major classes of factors cause sex differences (Arnold 2009; Arnold 2012; Arnold et al. 2013; Schaafsma and Pfaff 2014; Arnold 2017). First, some sex differences are caused by different circulating levels of ovarian and testicular hormones, known as “activational effects”. These differences are reversible because they are eliminated by gonadectomy of adults. Second, certain sex differences persist after gonadectomy in adulthood and represent the effects of permanent or differentiating effects of gonadal hormones, known as “organizational effects,” that form during development. A third class of sex differences are caused by the inequality of action of genes on the X and Y chromosomes in male (XY) and female (XX) cells, and are called “sex chromosome effects”.

To date, few studies have systematically evaluated the relative size and importance of each of these three classes of factors acting on phenotypic or global gene regulation systems (Arnold 2019). For instance, the activational effects of hormones have been established as a significant contributor to sexual dimorphism in metabolic diseases, with additional evidence pointing to sex chromosome effects on obesity and lipid metabolism (Chen et al. 2012; Link et al. 2015; Link et al. 2020). Previous studies have also emphasized the importance of organizational or activational hormone effects that contribute to sex differences in gene expression in the liver (Mode and Gustafsson 2006; van Nas et al. 2009; Waxman and Holloway 2009; Sugathan and Waxman 2013; Zheng et al. 2018). However, the tissue-specific contributions and the interactions of activational, organizational, and sex chromosome effects on gene regulation are poorly investigated.

Here we conduct a systematic investigation to understand the relative contribution of the three sex-biasing factors in gene regulation (**Figure 1)**. We used the Four Core Genotypes (FCG) mouse model, in which the type of gonad (ovary or testis) is independent of sex chromosome complement (XX or XY) (De Vries et al. 2002; Burgoyne and Arnold 2016). The model is a 2×2 comparison of the effects of sex chromosome complement by fixing the gonadal status (XX vs. XY with ovaries; XX vs. XY with testes) and type of gonad by fixing the sex chromosome type (ovaries vs. testes with XX genotype; ovaries vs testes with XY genotype). By varying adult gonadal hormone levels via gonadectomy and subsequent hormonal treatments we also asked how androgens and estrogens influence gene expression as a function of sex chromosome complement and gonadal sex. The design allows comparison of the magnitude of effect of each sex-biasing factor and the interactions among different factors.

**Figure 1.**
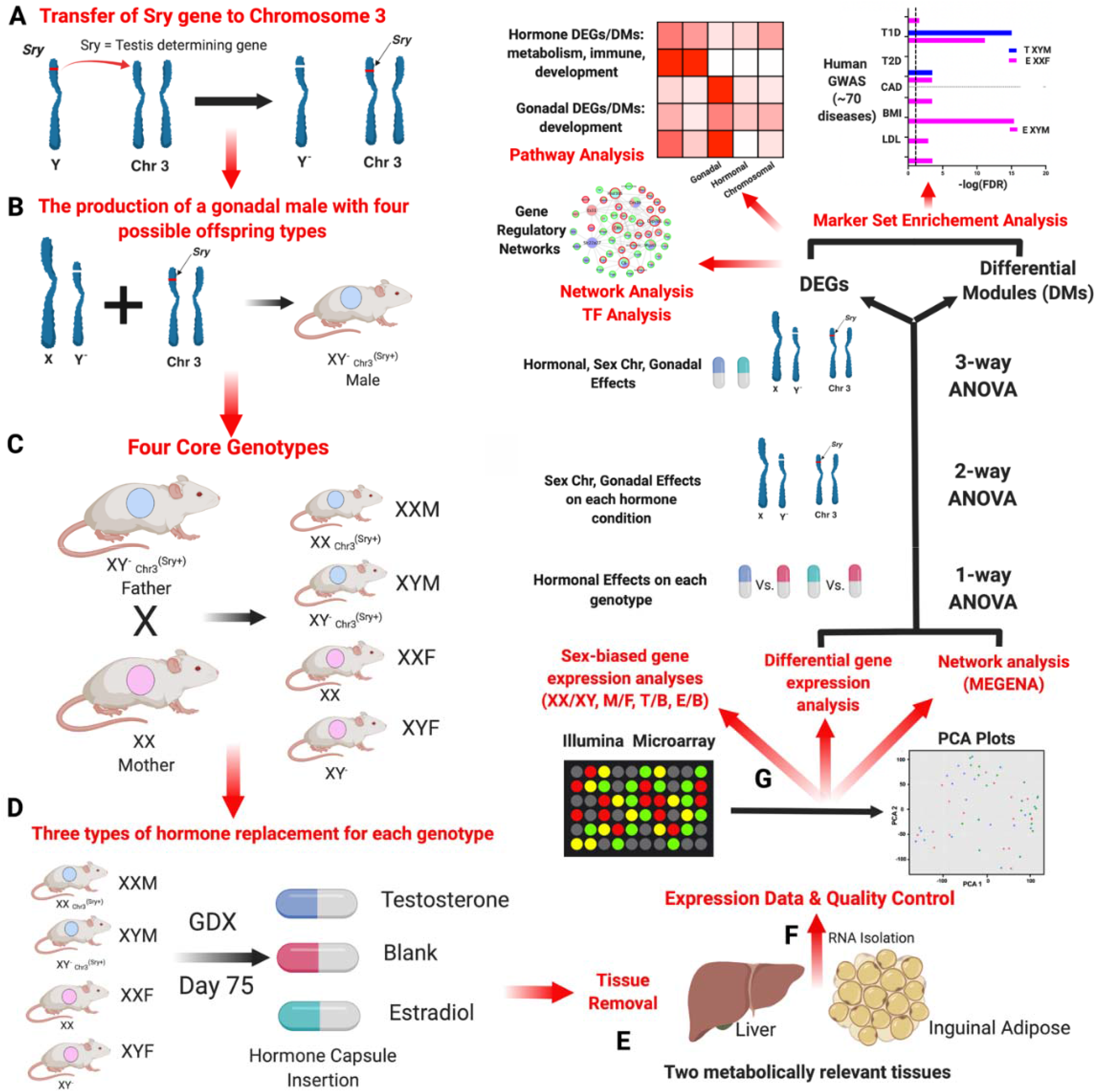
Overall Study Design. **A.** Transfer of the *Sry* gene to Chromosome 3. *Sry* which is usually located on the Y chromosome was deleted (a spontaneous deletion) and inserted as a transgene onto chromosome 3, making *Sry* independent of the Y chromosome. **B.** The production of a gonadal male XY^-^ Chr3^*Sry*+^, which has the ability to produce 4 types of gametes resulting in the four core genotypes (FCG). **C.** The generation of the FCG mice. Mating of XY^-^ _Chr3_^*Sry*+^ male and XX female produce four types of mouse offspring (two gonadal males and two gonadal females): XY^-^_Chr3_^*Sry*+^ (XYM), XX _Chr_^*Sry*+^ (XXM), XX (XXF), XY^-^ (XYF). **D.** Modulation of sex hormones in mouse offspring of each genotype after gonadectomy (GDX). Each of the four core genotypes underwent GDX at day 75 and was implanted with a capsule that contained either estradiol, testosterone or blank (n = 5/genotype/treatment). **E.** Dissection of the liver and inguinal adipose tissue for RNA isolation. **F.** Gene expression profiling and quality control. Using an Illumina microarray, we measured the transcriptome and then carried out a principal component analysis (PCA) to identify outliers and global patterns. **G.** Bioinformatics analyses. Differentially expressed genes (DEGs) influenced by individual sex-biasing factors were identified using 3-way ANOVA (chromosomal, gonadal and hormonal effects), 2-way ANOVA (gonadal and chromosomal effects under each hormone condition), and a 1-way ANOVA (estradiol and testosterone treatment effects in individual genotypes). Gene coexpression networks were constructed using MEGENA and differential coexpression modules (DMs) affected by individual sex-basing factors were identified using 3-way, 2-way, and 1-way ANOVAs. DEGs and DMs were analyzed for enrichment of functional categories or biological pathways. The relevance of the DEGs to human disease was assessed via integration with human genome-wide association studies (GWAS) for >70 diseases using the Marker Set Enrichment Analysis (MSEA). Transcription factor analysis and gene regulatory network analysis were additionally conducted on the DEGs derived from the oneway ANOVA.

Using the FCG model, our aim is to assess the role of the three sex-biasing factors on gene expression, molecular pathways, and gene network organization in the liver and adipose tissue, which are central tissues for metabolic and endocrine homeostasis, with adipose tissue additionally contributing to immune functions. We further aim to understand the influence or relationship of each sex-biasing factor in their contribution to various human diseases.

## RESULTS

### Overall study design

We used FCG mice on a C57BL/6J (B6) background. In FCG mice, the Y chromosome (from strain 129) has sustained a spontaneous deletion of *Sry,* and an *Sry* transgene is inserted onto chromosome 3 (Burgoyne and Arnold 2016). Here, “male” (M) refers to a mouse with testes, and “female” (F) refers to a mouse with ovaries. FCG mice include XX males (XXM) and females (XXF), and XY males (XYM) and females (XYF; **Figure 1**). A total of 60 FCG mice were gonadectomized (GDX) at 75 days of age and implanted immediately with medical grade Silastic capsules containing Silastic adhesive only (blank control; B) or testosterone (T) or estradiol (E). This study design produced 12 groups (**Figure 1**), with 4 groups of FCG mice (XXM, XXF, XYM, XYF) and each group subdivided into B or T or E based on hormonal treatment: XXM_B, XXM_T, XXM_E, XYM_B, XYM_T, XYM_E, XXF_B, XXF_T, XXF_E, XYF_B, XYF_T, XYF_E (n=5/genotype/treatment). Liver and inguinal adipose tissues were collected 3 weeks later for transcriptome analysis. All liver samples passed quality control (n=5/group) whereas 5 adipose samples across 4 of the 12 groups failed quality control (n=3-5/group; see Methods). The design allowed detection of differences caused by three factors contributing to sex differences in traits. (1) “Sex chromosome effects” were evaluated by comparing XX and XY groups (n=~30/sex chromosome type/tissue). (2) “Gonadal sex effects” were determined by comparing mice born with ovaries vs. testes (n=~30/gonad type/tissue). Since mice were analyzed as adults after removal of gonads, the gonadal sex effects represent organizational (long-lasting) effects of gonadal hormones, such as those occurring prenatally, postnatally, or during puberty. This group also includes effects of the *Sry* gene, which is present in all mice with testes and absent in those with ovaries. Any direct effects of *Sry* on non-gonadal target tissues would be grouped with effects of gonadal sex. (3) “Hormone treatment effects” refers to the effects of circulating gonadal hormones (activational effects) and were evaluated by comparing E vs. B groups for estradiol effects, and T vs. B groups for testosterone effects, with n=~20/hormone type/tissue.

### Global effects of sex chromosome complement, gonadal sex, and hormonal treatments on liver and adipose tissue gene expression

To visualize the overall gene expression trends due to effects of the three primary sex-biasing components, we conducted principal component analysis (PCA; **Supplemental Figure S1**). For the adipose tissue, hormonal treatment (**Supplemental Figure S1A**), sex chromosomes (**Supplemental Figure S1B**) and gonadal sex (**Supplemental Figure S1C**) did not clearly separate the groups. However, in the liver there was a separation of groups based on gonadal hormones, particularly in response to testosterone treatment (**Supplemental Figure S1D**), but not based on chromosomal or gonadal factors (**Supplemental Figure S1E, S1F**).

We then asked which individual genes in liver and adipose tissues were affected by adult hormone level, gonadal sex, and sex chromosome complement, as well as interactions between these factors, using three sets of ANOVA tests to address biological questions at different resolution. We defined a differentially expressed gene (DEG) as a gene that passed a false discovery rate (FDR)<0.05 for individual sex-biasing factors and the interaction terms from the ANOVAs. First, we used a 3-Way ANOVA (3WA) to test the main effects of sex hormones, gonad type, and sex chromosome as well as the interaction terms. Tens to thousands of DEGs were identified in liver (**Table 1**) and adipose tissue **(Table 2)**. In both tissues, hormonal treatments affected the largest numbers of genes, followed by fewer genes that were responsive to gonadal/organizational effects or sex chromosome complement (**Figure 2**). Testosterone treatment in the liver induced the largest number of DEGs (**Figure 2A**), whereas in inguinal adipose tissue estradiol treatment affected the greatest number of DEGs (**Figure 2D**). These trends remained when different statistical cutoffs (unadjusted p<0.05, p<0.01, FDR<0.1, FDR<0.05) were used (**Supplemental Figure 2**). These results support tissue-specific sensitivity to different hormones.

**Figure 2.**
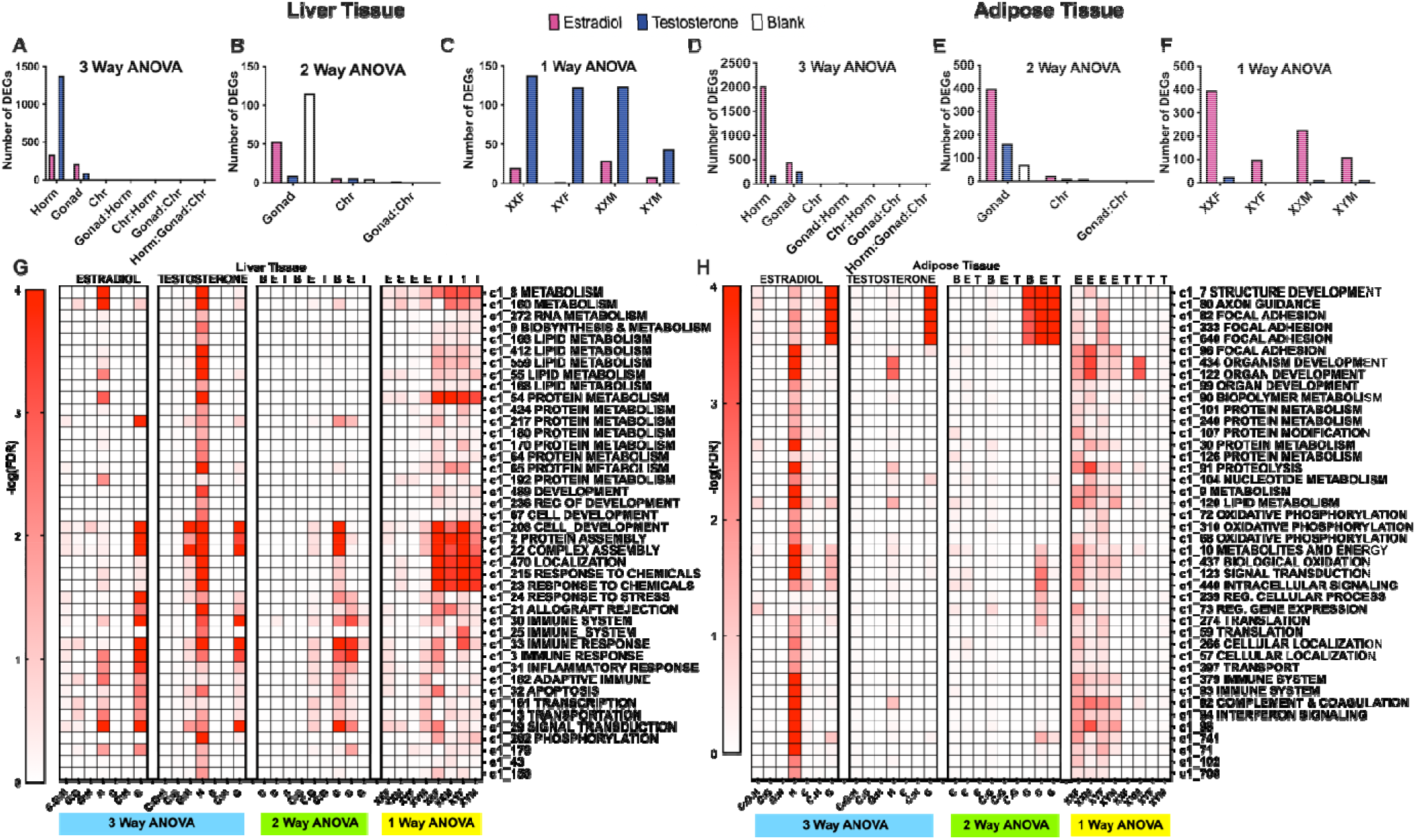
Bar graphs (A-F) and heatmaps (G-H) representing the number of DEGs for each sex-biasing factor and differential co-expression modules from a 3-way, 2-way, and 1-way ANOVA, respectively. Each bar graph represents the number of DEGs based on each specific statistical analysis at FDR<0.05. **A**, **D** represent results from 3-way ANOVAs run separately in testosterone vs. blank groups, and estradiol vs. blank groups to examine hormone, gonad, and sex chromosome effects as well as the interaction terms. **B** and **E** represent results from 2-way ANOVAs with factors of gonadal sex and sex chromosomes as well as the interaction term, run separately on data from testosterone (T), estradiol (E), and blank (B) treatment groups. **C** and **F** represent results from 1-way ANOVAs testing effects of hormonal treatments (vs. Blank) in each of the four genotypes for liver and inguinal adipose tissue. In A and D, bars are colored according to the hormonal treatment variable – data from estradiol vs. blank (pink), or data from testosterone vs. blank (blue). In panels **B** and **E**, colors represent the hormonal treatment condition (testosterone groups blue, estradiol groups pink, and blank groups white) for each of the separate 2-way ANOVAs. In panels **C** and **F**, colors show effects of testosterone vs blank (blue) or estradiol vs blank (pink) in each of the four genotypes. Horm = Hormone, Chr = Sex Chromosome, M = Testes/Sry present, F = Ovaries present, no Sry. **G** represents the heatmap for liver. **H** represents the heatmap for adipose tissue. Each heatmap shows results from 1-way ANOVA, 2-way ANOVA and 3-way ANOVA for each test factor hormone (H), chromosome (C), and gonad (G) when treated with testosterone (T), estradiol (E) and blank (B). Interaction terms among H, C, and G were also tested. For instance, C:G:H indicates the interaction term among the 3 factors in 3-way ANOVA. The influence of each sex-biasing factor on the coexpression modules was assessed using the first principal component of each module to represent the expression of that module, followed by 3WA, 2WA, 1WA tests and FDR calculation to identify differential modules (DMs) at FDR <0.05 that are influenced by the various sex-biasing factors. Each module was annotated with canonical pathways from GO and KEGG. When a module is not annotated with a pathway name, it did not show significant enrichment for genes in any given pathway tested. Colors correspond to the statistical significance of the effects of sex factors on modules in the form of −log10(FDR).

**Table 1:** Liver DEGs affected by sex-biasing factors and the associated GO/KEGG pathways. Only DEGs with main effects are shown; full DEG lists (Supplemental Table S1) and DEGs significant from analysis of interaction terms between sex-biasing factors are in Supplemental Table S2.

| Analysis | Treatment/<br>Genotype<br>group | Sex<br>Factor | DEGs at<br>FDR <<br>0.05 | Top Five DEGs<br>Based on FDR | Top DEGs<br>Based on LogFC | Top GO/KEGG Pathways<br>FDR<0.05 |
| --- | --- | --- | --- | --- | --- | --- |
| **3-way**<br>**ANOVA** | **T + B** | Horm | **1378** | *Trim47l, Cux2, Elovl3, Cyp3a41, Sult3a1* | **Up:** *Elovl3, Serpina4-ps1, Cyp4a12, Cyp2u1, Slco1a1*<br>**Down:** *Cyp2b13, Fmo3, Cux2, Trim47l, Eci3* | Lipid metabolic process<br>Organic acid and metabolic process<br>Cellular lipid metabolic process<br>Primary bile acid biosynthesis |
|  |  | Gonad | **93** | *Cyp3a41, Sult3a1, Cxcl9, Siat10, Ly6a* | **Up:** *Cml4, Cyp4a12, Asns, Lcn13, Cpne8*<br>**Down:** *Gbp1, BC013712, Spic, Sult3a1, Cyp3a41* | Defense response<br>Immune system process<br>Response to stress |
|  |  | Chr | **8** | *Ddx3y, Eif2s3y, Xist, Kdm6a, Tmsb4x* | **Up:** *Eif2s3y, Ddx3y, Tmsb4x*<br>**Down:** *Xist, Ntrk2, Kdm6a, Eif2s3x, Wfdc2* | Rig I like receptor signalling pathway<br>Transmembrane receptor protein tyrosine kinase signalling pathway<br>MAPK signaling pathway |
|  | **E + B** | Horm | **333** | *Cyp17a1, Slc11a1, Trp53inp2, Clqb, Clmn* | **Up:** *Prtn3, Lcn13, Isynal, Ear4, Acot11*<br>**Down:** *Adam11, Gsta1, 1700009P17Rik, Pde6c, Tppp* | Carboxylic acid metabolic process<br>Organic acid metabolic process<br>Monocarboxylic acid metabolic process<br>Complement and coagulation cascades |
|  |  | Gonad | **209** | *Cxcl9, Cyp3a41, Ii, Slc11a1, Ccr5* | **Up:** *Cml4, Cyp4a12, Tiam2, Susd4, Agpat6*<br>**Down:** *Cyp3a41, Thy1, BC013712, Tbc1d10c, Cyp4f18* | Defense response<br>Lipid metabolic process<br>Immune response (SLE) |
|  |  | Chr | **8** | *Eif2s3y, Xist, Ddx3y, Kdm6a, Eif2s3x* | **Up:** *Eif2s3y, Hccs, Ddx3y, Tmsb4x, Mgm1*<br>**Down:** *Xist, Kdm6a, Eif2s3x* | Electron transport<br>Ubiquitin mediated proteolysis<br>Organ morphogenesis |
| **2-way**<br>**ANOVA** | **T** | Gonad | **9** | *Lcn13, Cml4, Siat10, Ii, Cxcl9* | **Up:** *Asns, Lcn13, Cml4, Cyp4a12, Gm4956*<br>**Down:** *Li, Cxcl9, Siat10, Qpct* | Cellular response to stress/nutrient levels/extracellular stimulus<br>Amino Acid Synthesis |
|  |  | Chr | **6** | *Ddx3y, Eif2s3y, Xist, Kdm6a, Tlr7* | **Up:** *Eif2s3y, Tlr7, Ddx3y, Tmsb4x*<br>**Down:** *Kdm6a, Xist* | Traf6 mediated irf7 activation in Tlr7 signaling<br>Interleukin 8 biosynthetic process |
|  | **E** | Gonad | **53** | *H2-Ab1, H2-Eb1, H2-DMb1, Cd44, Ii* | **Up:** *Cyp7b1, Plekhh1, Actr1b, Sh3glb2*<br>**Down:** *Thy1, Tbc1d10c, Cd79b, A1467606, S100a8* | Asthma<br>Leishmania Infection<br>Allograft rejection<br>Systemic Lupus Erythematosus |
|  |  | Chr | **6** | *Eif2s3y, Xist, Ddx3y, Als2cl, Kdm6a* | **Up:** *Eif2s3y, Als2cl, Ddx3y*<br>**Down:** *H2-DMb1, Kdm6a, Xist* | Asthma<br>Allograft rejection |
|  | **B** | Gonad | **115** | *Cyp3a41, Sult3a1, Slc11a1, Susd4, Nox4* | **Up:** *Cml4, Cyp4a12, Cyp2d9, Igfbp2, Susd4*<br>**Down:** *Sult3a1, Cyp3a41, A1bg, C1orf38, Spic* | Systemic Lupus Erythematosus<br>Functionalization of Compounds<br>CYP450 arranged by substrate type<br>Biological oxidations |
|  |  | Chr | **5** | *Eif2s3y, Xist, Ddx3y, Kdm6a, Ntrk2* | **Up:** *Eif2s3f, Ddx3y*<br>**Down:** *Xist, Ntrk2, Kdm6a* | Rig I like receptor signaling pathway<br>Aging, Circadian Rhythm<br>Fatty Acid Metabolism |
| **1-way**<br>**ANOVA**<br>**Post-hoc** | **XXF** | T | **139** | *Aqp4, A1bg, Gas6, Parp1* | **Up:** *Cyp4a12b, Serpina4-ps1, Elovl3, Cyp2u1, Aqp4*<br>**Down:** *A1bg, Cyp2b13, Fmo3, Slc22a26, Cux2* | Biological Oxidations<br>Metabolism (xenobiotic/drug/glutathione)<br>Immune (complement/interferon) |
|  | **XYF** | T | **123** | *Heg1, Aqp4, Cyp2u1, I11a, Fam111a* | **Up:** *Cyp4a12, Elovl3, EG241041, Cyp2u1, Fst*<br>**Down:** *Cyp2b13, Fmo3, A1bg, Cux2, Eci3* | Biological Oxidations<br>Complement cascade<br>Bile acid metabolism<br>Metabolism (steroid/drug) |
|  | **XXM** | T | **124** | *Spry4, Cutl2, Sybu, Akr1d1, Slco1a1* | **Up:** *Serpina4-ps1, Elovl3, Lcn13, Slco1a1, Cyp4a12a*<br>**Down:** *Cux2, Eci3, Trim47l, Irx3, Akr1d1* | Biological Oxidations<br>Immune system (complement/SLE)<br>Cell development/Wnt signaling<br>Drug/Protein/Bile acid/salt metabolism |
|  | **XYM** | T | **44** | *Atp2a3, Unc79, Trim47l, Slco1a1, Lcn13* | **Up:** *Serpina4-ps1, Elovl3, Slco1a1, Lcn13, Gadd45g*<br>**Down:** *Fmo3, Cux2, Slc22a26, Trim47l, Unc79* | Cell development / Localization<br>Metabolism (Drug/Riboflavin)<br>Immune (Complement/Innate) |
|  | **XXF** | E | **20** | *Zfp367, Lamb2, Ahcy12, Prtn3, Pdlim4* | **Up:** *Prtn3, Clqb, Li, Clqc, Visg4*<br>**Down:** *Ahcy12, Lamb2, Zfp367* | Signal transduction<br>Metabolic Pathways<br>Complement Pathways<br>Class B2 Secretin Family Receptors |
|  | **XYF** | E | **2** | *Klk1b27, Reck* | **Down Only:** *Klk1b27, Reck* | Developmental Processes |
|  | **XXM** | E | **29** | *Slco1a, Tasor2, Ywhae, Sult3a1, Lcn13* | **Up:** *Sult3a1, Lcn13, Slco1a1, Slc11a1, Serpina1e*<br>**Down:** *Ywhae, BC016423, Ankrd12, Spc24, Serpina6* | Nitrogen/Vitamin/ Steroid Hormones<br>Metabolism<br>Complement/Inflammatory Pathways<br>Signal Transduction/Transcription |
|  | **XYM** | E | **8** | *Adam11, Sult3a1, Spc24, Lfit2, Gsta4* | **Up Only:** *Lfit2, Sult3a1, Cyp17a1, Gsta4, Serpina6* | Immune response (interferon signaling)<br>Protein complex assembly<br>Biological Oxidations/ Metabolism of steroid hormones/Vitamins |

**Table 2:** Inguinal adipose DEGs affected by sex-biasing factors and the associated GO/KEGG pathways. Full DEG lists (Supplemental Table S3) and DEGs significant from analysis of interaction terms between sex-biasing factors are in Supplemental Table S2.

| Analysis | Treatment/<br>Genotype<br>Group | Factor | DEGs at<br>FDR <<br>0.05 | Top DEGs Based on<br>FDR | Top DEGs Based on LogFC | GO/KEGG Pathways at FDR<0.05 |
| --- | --- | --- | --- | --- | --- | --- |
| **3-way**<br>**ANOVA** | **T + B** | Horm | **275** | *Mpp5, Hp, Lrg1, Serpina3n, Prtn3* | **Up:** *Serpina3m, Mt2, Agt, Gm1157, Krt36*<br>**Down:** *Fabp5, Cartac1, Clca5, Rad54b, P2rx5* | Cell Cell Adhesion<br>Protein Signaling Pathway |
|  |  | Gonad | **268** | *Mpp5, Dnaic1, Aida, Cited1, Slc12a2* | **Up:** *Rab9b, Sntg2, Dusp15, Pinx1, 1100001G20Rik*<br>**Down:** *Dnaic, Aida, Nrnm3, Rab25, Upk2* | Multicellular Organismal Development<br>Cell Cell Adhesion<br>Anatomical Structure Development<br>Pathways in Cancer |
|  |  | Chr | **15** | *Mpp5, Xist, Ddx3y, Eif2s3y, Hccs* | **Up:** *Eif2s3y, Jarid1d, Ddx3y, Mpp5, Tlr7*<br>**Down:** *Xist, Aatk, Kdm6a, Eif2s3x, Il11ra1* | Interleukin 8 biosynthetic process |
|  | **E + B** | Horm | **2029** | *Dnaic1, Gas6, Greb1, Fbxw17, Lrg1* | **Up:** *Adamts19, Hpcra, Rpp25, Greb1, Thbs1*<br>**Down:** *F2rl3, Fabp5, Clca5, Crtac1, Ucp1* | Protein Metabolic Process<br>Respiratory Electron Transport<br>Focal Adhesion |
|  |  | Gonad | **449** | *Dnaic1, Prr1, Prr15l, Atad4, Ercc2* | **Up:** *Dusp15, Mtap7d3, Rab9b, Gm525, Kcnh2*<br>**Down:** *Prr15l, Atad4, Ercc2, Cldn3, Prom2* | Anatomical Structure Development<br>Nitrogen Compound Metabolic Process<br>Aldosterone Sodium Reabsorption |
|  |  | Chr | **10** | *Xist, Dnaic1, Ddx3y, Eif2s3y, Eif2s3x, Hcc* | **Up:** *Eif2s3y, Ddx3y, Tlr7, Hccs, Gprasp1*<br>**Down:** *Xist, Gm525* | Rig I like receptor signaling pathway<br>Generation of precursor metabolites and energy |
| **2-way**<br>**ANOVA** | **T** | Gonad | **161** | *Rbm35a, Sephs1, Ferm5l, Wnt7b, Lrrc26* | **Up:** *Rab9b, Mlec, Por, Cdsn, Rprml*<br>**Down:** *BC004852, Wnt7b, Al646023, Rbm35a, Fermt1* | Epithelial Cell Differentiation<br>Excretion<br>Cell Cell Communication<br>Cell Carcinoma/ Junction Organization |
|  |  | Chr | **11** | *Eif2s3y, Rbm35a, Sephs1, Fermt1, Ddx3y* | **Up:** *Eif2s3y, Ddx3y, Wnt7b, Fermt1, BC004853*<br>**Down:** *Xist, Kdm6a* | Seleno amino acid metabolism<br>Electron transport<br>Basal Cell Carcinoma |
|  | **E** | Gonad | **400** | *Mtap7d3, Elf3, Ercc2, Pkp1, Slc5a7* | **Up:** *Kcne11, Dusp15, Kcnh2, Mtap7d3, Gm525*<br>**Down:** *Pkp1, Fermt1, Atad4, Elf5, Mfsd2a* | Arginine & Proline Metabolism<br>Cell Junction Organization<br>Cell Cell Communication<br>Epidermis Development |
|  |  | Chr | **22** | *Mtap7d3, Xist, Eif2s3y, Ddx3y, Elf3* | **Up:** *Eif2s3y, Ddx3y, Tlr7, Hccs, Mxra7*<br>**Down:** *Xist, Mtap7d3, Kdm6a, Fermt1, Eif2s3x* | Neurotransmitter release<br>Cell Differentiation |
|  | **B** | Gonad | **70** | *Rnf208, Cited1, Dnaic1, Prr15l, Mpp5* | **Up:** *Cln3, Mpp5, Bikl, Adck5, Capn5*<br>**Down:** *Prr15l, Dnaic1, Cited1, Rnf208, Tcfap2b* | Cell Cell Communication<br>YAP1 and WWTR1 TAZ stimulated gene expression<br>Epidermis Development |
|  |  | Chr | **10** | *Xist, Mpp5, Rnf208, Cited1, Dnaic1* | **Up:** *Mpp5, Ddx3y, Rnf208, Prr15l, Cited1*<br>**Down:** *Xist, Eif2sx, Kdm6a* | Tight Junction interactions<br>Electron Transport<br>Rig I like receptor signaling |
| **1-way**<br>**ANOVA**<br>**post-hoc** | **XXF** | T | **26** | *Rad9b, Rfwd3, BC049762, Dusp15, Crtac1* | **Up:** *Slc47a1, Pitpnc1, Mpp5, sc10003251.1_21, Rfwd3*<br>**Down:** *H2-Q10, Crtac1, Slc4a1, Dusp15, Rad9b* | Cell Carcinoma<br>Cell Development/Differentiation/Wnt Signaling<br>Immune (Antigen Presentation) |
|  | **XYF** | T | **1** | *Fabp5* | **Down Only:** *Fabp5* | Development Processes |
|  | **XXM** | T | **13** | *P2rx5, Krt36, Grik5, F2rl3, AA792892* | **Up:** *Mt2, Krt36, Hp, Serpina3n, Lrg1*<br>**Down:** *Aspg, Clca5, F2rl3, Grik5, AA792892* | Organ Development/Cell Division/cycle<br>Immune (haemostasis, antigen presentation) |
|  | **XYM** | T | **13** | *Sephs1, Col4a5, Abp1, Tnnt1* | **Up:** *Mt2, Apom, Tnnt1, Fam25c, Col4a5*<br>**Down:** *Cxcl15, Abp1, Sephs1, Rnf144a* | Cell proliferation/Migration |
|  | **XXF** | E | **395** | *Echdc1, Spin, Il10, Rad9b, S100a14* | **Up:** *Ppp1r27, Adamts19, Saa3, Thbs1, Krt15*<br>**Down:** *Dnaic1, Cited1, Aida, Fam84a, Clca5* | Immune System (Interferon, Antigen presentation)/ ERK pathways<br>Metabolism (Porphyrins, tryptophan)<br>Cancer/Protein Stabilization |
|  | **XYF** | E | **100** | *Fam83f, Csf3r, Galnt9, S100a14, Efna4* | **Up:** *Tph1, Areg, Abp1, Ly6g6c, Lgals7*<br>**Down:** *Actc1, Aqp4, Gdap1, A930018M24Rik, Dnaic1* | Complement Cascade<br>Transport by Aquaporins<br>Development (Cell migration, cell junction organization, ECM R interaction, Focal Adhesion) |
|  | **XXM** | E | **228** | *Al646023, Eraf, P2rx5, Ildr1, 6330403K07Rik* | **Up:** *Ifi205, Hpcra, Rpp25, Il1rl1, Nxn12*<br>**Down:** *Anxa8, Clca5, F2rl3, Grik5, Grb14* | Complement Cascade/Interferon Signaling<br>ECM Organization<br>Metabolism (glucose, glutathione, lipid) |
|  | **XYM** | E | **110** | *1700088E04Rik, 6330403K07Rik, Dlx3, 1110012J17Rik, Cacna1g* | **Up:** *Hpcra, Spon1, Rpp25, Thbs1, Il31ra*<br>**Down:** *Fabp5, Ccno, Trfr2, F2rl3, Cidea* | Immune (Interferon, Tlr4/ERK signaling)<br>Metabolism (Nicotinamide, Glutathione, lipid)/ EGF Pathway/Cell Proliferation |

Next, we asked if the sex chromosome and gonadal effects are more evident in specific hormonal treatment groups using a 2-way ANOVA (2WA). In the liver, the organizational effects of gonad type were strongest in gonadectomized mice without hormone replacement (blank group) (**Figure 2B**). By contrast, in adipose tissue the gonadal sex effect was most prominent in the estradiol treated groups (**Figure 2E**), suggesting that estradiol levels augment the enduring differential effects of gonads on the adipose tissue transcriptome. Sex chromosome effects were limited regardless of hormonal treatment status, but captured specific genes important in inflammatory signaling and metabolic pathways.

Lastly, we examined whether the effects of testosterone and estradiol are dependent on genotypes using a 1-way ANOVA (1WA) followed by post-hoc analysis. More liver genes were affected by testosterone than by estradiol regardless of genotype, although XYM liver appeared to be less responsive to testosterone than liver from other genotypes (**Figure 2C**). By contrast, in adipose tissue, estradiol affected more DEGs in XX genotypes (XXM and XXF) than in XY genotypes (XYM and XYF), whereas testosterone had minimal impact on adipose tissue gene expression in all four genotypes (**Figure 2F**). These results further support tissue-specific effects of estradiol in adipose tissue and testosterone in liver, and indicate that activational effects of hormones also depend on sex chromosome complement and hormonal history (gonadal sex) of the animal.

### Genes and pathways affected by hormonal treatment

In the liver, the 3WA analysis showed that testosterone treatment induced the greatest number of DEGs with 1378 compared to 333 DEGs from estradiol treatment (**Table 1; Figure 2A)**. The testosterone DEGs were enriched for numerous metabolic pathways including lipid metabolism, organic acid metabolism, and bile acid biosynthesis (**Table 1**). Within individual genotypes, testosterone treatment altered 123-139 DEGs in the XXF, XYF, XXM genotypes but only 44 DEGs in the XYM genotype (post-hoc 1WA in **Table 1**), with development, response to chemicals, immune response, and metabolism being the key over-represented terms. The 333 liver DEGs from estradiol treatment in the 3WA analysis showed enrichment for metabolic (organic acid metabolism, carboxylic acid metabolism) and immune pathways (complement and coagulation). The post-hoc 1WA for estradiol versus blank treatment in individual genotypes showed a high of 29 DEGs for XXM and a low of 2 DEGs for XYF (**Table 1**), again with metabolism and immune pathways showing enrichment.

In contrast to liver, we found that the effect of estradiol treatment was more profound (2029 DEGs) than that of testosterone (275 DEGs) in 3WA of the inguinal adipose tissue (**Table 2; Figure 2D**). These DEGs were enriched for protein metabolism, focal adhesion, and transport pathways. The post-hoc 1WA of estradiol effect showed 395 DEGs for XXF, while XXM, XYF, and XYM each had more than 100 DEGs (**Table 2; Figure 2F**), with enrichment for developmental, immune, and lipid metabolic processes. Testosterone treatment produced 275 DEGs in adipose tissue in 3WA and a few to tens of DEGs in individual genotypes in 1WA. Pathways enriched include cell-cell adhesion, development, regulation of transcription, and protein signaling.

Overall, both estradiol and testosterone affected genes involved in metabolism, development, and immune function. However, estradiol primarily affected these processes in the adipose tissue, whereas testosterone exhibited influence in the liver.

### Genes and pathways affected by gonadal sex

In the liver, 3WA analyses revealed 93 DEGs influenced by gonadal sex when testosterone and blank treatment groups were considered, and 209 DEGs in the analysis of estradiol and blank groups (**Table 1**). These genes were enriched for immune/defense response and lipid metabolism pathways. To further tease apart the effect of gonadal sex in individual hormonal treatment groups, we conducted a 2WA and found that gonadal sex has the strongest influence on inflammatory and metabolism genes in the absence of hormones (blank group; 115 DEGs), whereas the gonadal sex effects were reduced by estradiol treatment (53 DEGs) and minimized in the testosterone group (9 DEGs; **Table 1**; **Figure 2B**).

For the inguinal adipose tissue, gonadal sex showed strong effects on gene expression, with more than twice as many DEGs as in liver tissue in the 3WA analysis (**Figure 2A** vs. **2D**). Genes involved in developmental processes were enriched in these DEGs. Further dissection of the gonadal sex effect in individual hormonal treatment groups in a 2WA analysis showed that the effects of gonadal sex were strongest in the estradiol treatment group (400 DEGs), followed by testosterone group (161 DEGs), and lastly by the blank group (70 DEGs; **Figure 2E**). In addition to affecting development related genes, gonadal sex also affected genes involved in arginine and proline metabolism in the estradiol treatment group, and cancer-related genes in the testosterone treatment group.

These results support the importance of gonadal sex in regulating development processes in both tissues. However, in the liver, hormonal treatments minimized the effects of gonadal regulation of gene expression, whereas in the adipose tissue, hormones amplified the gonadal influence on gene expression to regulate additional metabolism genes (by estradiol) or cancer-related genes (by testosterone). In both tissues, the gonadal sex effect was more prominent in the estradiol-treated group than in the testosterone-treated group (**Figure 2B** vs. **2E**).

### Genes and pathways affected by sex chromosome complement

In both the 3WA and 2WA analyses, ten or fewer genes were found to be significantly affected by sex chromosome complement at FDR < 0.05 in the liver (**Table 1; Figure 2C**) and 10-22 DEGs were influenced by sex chromosomes in the adipose tissue (**Table 2; Figure 2F**). Not surprisingly, these genes were mainly sex chromosome genes known to exhibit sex differences in gene expression levels, including *Xist*, *Ddx3y*, *Kdm6a*, *Hccs*, *Cited1*, *Tlr7* and *Eif2s3x/y* (Chen et al. 2012; Berletch et al. 2015; Golden et al. 2019; Itoh et al. 2019). However, autosomal genes were also influenced by sex chromosome type in both liver (e.g., *Ntrk2* and *H2-dmb1*) and adipose tissue (e.g., *Mpp5*, *Rbm35a*, and *Dnaic1*). Overall, the genes influenced by sex chromosome complement are involved in diverse processes including inflammation/immune response (*Tlr7*, *H2-dmb1*, *Cited1*), GPCR signaling (*Rbm35a*), metabolism (*Hccs*), and cell junction organization (*Mpp5*).

### Genes and pathways affected by interactions of sex-biasing factors

The interactions among the sex-biasing factors are supported by numerous DEGs with significant effects from the interaction terms in the ANOVA analyses (FDR<0.05; **Supplemental Table S2**). For instance, in adipose tissue 31 DEGs were affected by interactions between estradiol and gonad type. These DEGs were enriched in pathways such as VLDL particle assembly and regulation of leukocyte chemotaxis. DEGs *Dnaic1* and *Cited1* were expressed in female gonads (XXF or XYF) when no sex hormones were provided; genes such as *Ctns*, *Slc2a3*, *S100a14* and *Ler3* showed a significant increase in expression when estradiol treatment was provided to female gonads (**Supplemental Figure S3**). In the liver fewer genes showed significant interaction effects between pairs of sex-biasing factors (FDR<0.05) (**Supplemental Table S2**). For instance, expression of *Cyp3a41*, *Sult3a1* and *Cyp17a1* is downregulated by testosterone in mice with female gonads; *Lcn13* expression is upregulated by testosterone in mice with male gonads; expression of *Igfbp2* is upregulated by testosterone on female gonads but downregulated by testosterone on male gonads (**Supplemental Figure S4**).

### Comparison of mouse DEGs affected by sex-biasing factors with human sex-biased genes

To cross-validate the DEGs identified in our FCG mouse model, we compared them with sex-biased genes identified in human GTEx studies of liver (**Supplemental Table S4**) and adipose tissues (**Supplemental Table S5**) (Oliva et al. 2020). We found 29 out of 500 sex-biased genes (5.8%) in GTEx liver and 54 out of 500 sex-biased genes (10.8%) in GTEx adipose tissue were identified as DEGs affected by one or more sex-biasing factors in our FCG model. It is important to note the key difference between studies: the sex-biased genes in GTEx are the results of the combined effects of all sex-biasing factors whereas our FCG mouse study focuses on the effect of individual sex-biasing factors.

As GTEx studies cannot isolate specific sex biasing factors, our FCG model suggest hypotheses regarding the particular factors contributing to the sex biased genes found in humans. For instance, in adipose tissue, the GTEx female biased genes *ASAH1*, *PRDX2* and *LOXL1* might be explained by an effect of estradiol. In contrast, the male-biased adipose gene *HSD11B1* in GTEx can be explained in the FCG by the effect of testosterone (**Supplemental Figure S5**). In the liver, the human male-biased genes *ADH4*, *GNA12*, *HSD17B12* can be explained in our mouse model by an effect of testosterone, whereas the female-biased human genes *AS3MT*, *ZFX* and *CXCL16* were found to be affected by estradiol in FCG mice (**Supplemental Figure S6**). Therefore, the FCG mice not only can recapitulate sex-biased genes in human studies but suggest the specific sex-biasing factors that contribute to the sex bias.

### Coexpression modules affected by each sex-biasing factor

The above DEG analyses focused on genes that were individually influenced by sex-biasing factors as well as their interactions. Sets of genes that are highly coregulated or co-expressed can offer complementary information on coordinated gene regulation by sex-biasing factors that might be missed by the DEG-based analyses. To this end, we constructed gene coexpression networks for each tissue using MEGENA and identified 326 liver and 131 adipose coexpression modules. The first PCs of the coexpression modules, each composed of coregulated genes, were assessed for influence by sex chromosome, gonadal sex, and hormonal treatment factors using 3-, 2-, and 1-WAs.

In the heatmaps summarizing the significance of impact of each sex-biasing factor on individual modules (**Figure 2G, 2H**), we confirmed the large effect of hormonal treatment in regulating modules enriched for diverse biological pathways. In the liver, testosterone affected modules involved in metabolism (RNA, lipid, protein), development, protein assembly, chemical response, immune system (inflammation, adaptive immune response), apoptosis, and transcription/translation. In adipose tissue, estradiol influenced modules related to focal adhesion, development, metabolism (protein, lipid, oxidative phosphorylation), immune system (complement and coagulation), and translation.

Gonadal sex also showed considerable influence on liver modules related to protein metabolism/assembly, development, stress/immune response, apoptosis, and transcription/translation regulation, whereas in adipose tissue gonadal sex mainly affected developmental and focal adhesion processes, and to a lesser degree, lipid metabolism, biological oxidation, and intracellular signaling modules (**Figure 2G**).

The coexpression network analysis also confirmed the limited effect of sex chromosomal variation on altering coexpression modules (**Figure 2G, 2H**). However, in adipose tissue, sex chromosomes showed weak effects on modules related to lipid metabolism and intracellular signaling when estradiol and blank groups were considered, but not when the testosterone group was included **(Figure 2H)**.

Overall, the gene coexpression network analysis offered clearer patterns of tissue-specificity and functional specificity of each sex-biasing factor compared to the DEG-based analysis.

### Bulk tissue deconvolution to understand cellular composition changes through sex-biasing factors

To explore whether the DEGs and pathways/modules identified in FCG can be explained by cellular composition changes affected by each sex biasing factor, we carried out cell composition deconvolution analysis on the bulk tissue transcriptome data using CIBERSORTx based on single cell reference datasets of the corresponding tissues (see Methods). We subsequently assessed the hormonal, gonadal, and sex chromosomal effects on individual cell types.

In both the liver (**Supplemental Figure S7**) and adipose tissue (**Supplemental Figure S8**), hormones affected the largest number of cell types in terms of their abundance, including various immune cell populations such as the hepatocellular stellate cells (HSCs) and neutrophils in the liver, and macrophages, CD4 T-cells, dendritic cells, and antigen presenting cells in adipose tissue. Hormones also affected dividing cell populations and endothelial cells in both tissues. These cell populations affected by hormones support the DEGs and pathways involved in immune functions and development that are influenced by the same sex-biasing factor. Similar to the findings based on DEG and pathways analysis, the gonadal effect on cell populations is also dependent on the tissue and other sex-biasing factors: female gonads exhibit increases in hepatocytes, endothelial, and HSCs in the liver on an XX background, whereas male gonads showed an increase in macrophage proportion in adipose tissue on an XY and testosterone background. Lastly, the sex chromosome effect can be noted in immune cell populations, but it is generally dependent on the interactions with other sex factors. Overall, the changes in cellular composition support the changes in the pathways highlighted through our DEGs including immune, developmental and metabolic signals in both tissues.

### Effect of hormonal treatment on gene expression direction across genotypes

Due to the dominant effect of hormonal treatment as compared to gonadal sex or sex chromosome differences based on the above analyses, we further investigated the differences between testosterone and estradiol treatments in terms of the gene sets they target and the direction of gene expression change within and between tissues.

#### Overlapping DEGs between testosterone and estradiol treatment

Comparing groups of DEGs regulated by testosterone or estradiol in the 3WA (**Supplemental Figure S9**), 226 overlapped in the liver and 383 overlapped for adipose tissue. However, estradiol DEGs in individual genotypes did not overlap with those caused by testosterone in 1WA (p<0.05) in any genotype group (**Figure 3**). In particular, for the XYF mouse we found no overlapping DEGs in either the liver or adipose DEGs between testosterone and estradiol (**Supplemental Figure S10**). For other genotypes, we found an insignificant overlap of DEGs affected by the two hormones in the liver (**Figure 3A; 3B; 3C**) and adipose tissues (**Figure 3D; 3E; 3F**), and the direction of gene expression changes generally agreed between tissues. However, there were a few exceptions. *Fmo3* (Flavin containing monooxygenase 3, important for the breakdown of nitrogen-containing compound) in XXM liver (**Figure 3A**), and all the shared DEGs in XXF liver (*C1qb*, *C1qc*, and *Vsig4*; complement pathway genes) (**Figure 3C**) were affected by testosterone (down) and estradiol (up) oppositely.

**Figure 3.**
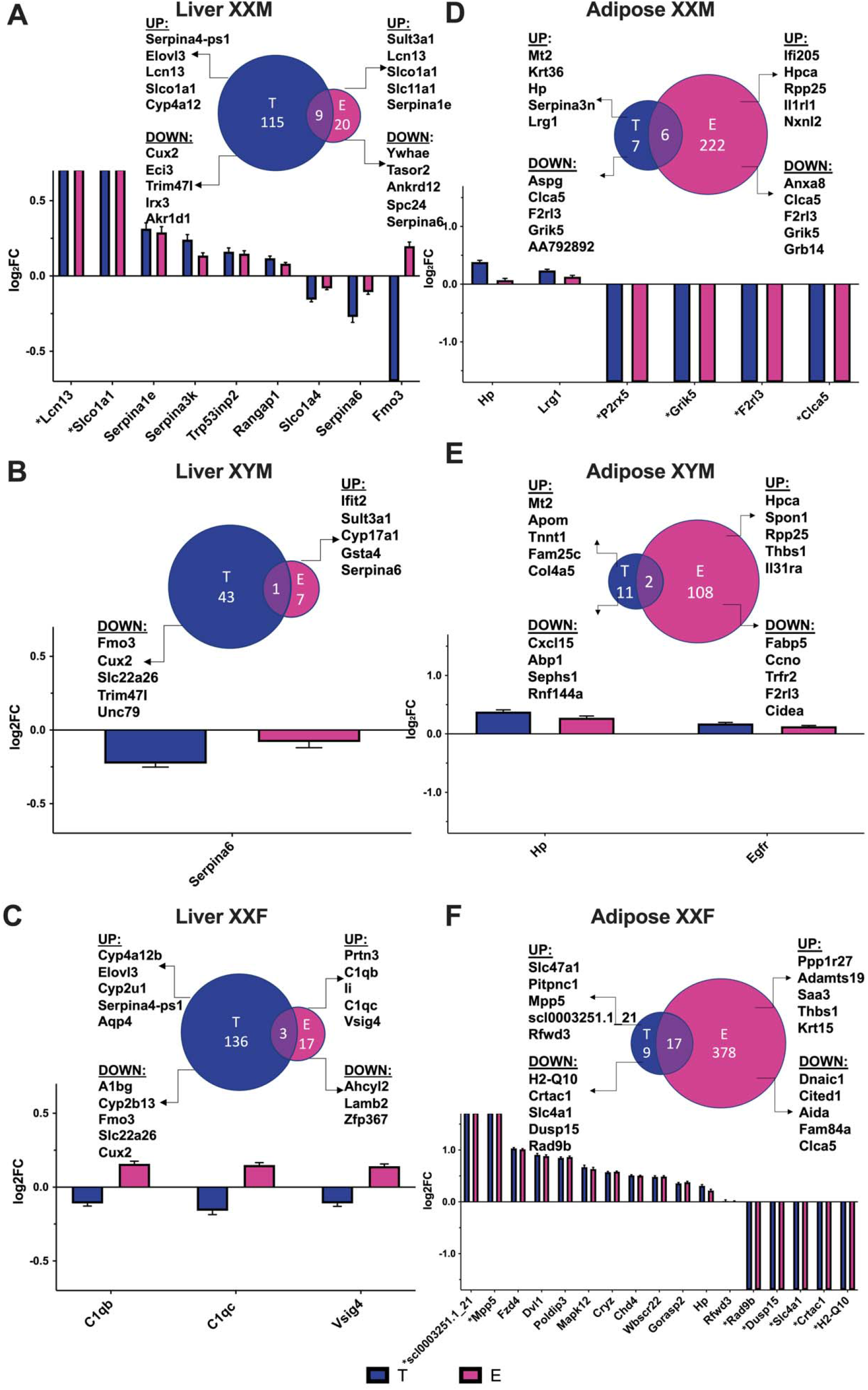
Venn Diagrams of DEG comparisons and Bar Graphs of overlapping DEGs between estradiol (E vs blank, abbreviated as E) and testosterone (T vs blank, abbreviated as T) treatment for each genotype in liver and adipose. **A**. Liver XXM. **B**. Liver XYM. **C**. Liver XXF. **D**. Adipose XXM. **E**. Adipose XYM. **F**. Adipose XXF. The bar graphs focused on the DEGs that passed an FDR < 0.05 and were overlapping between testosterone and estradiol treatment for each genotype and tissue. To understand the effects of each hormone, we plotted the log2 fold change of the hormonal effects. The Venn diagrams showcase comparison of DEGs of T effect vs E effect, as well as the top 5 up and down regulated genes for T or E in liver or adipose tissue for each genotype. There was no statistically significant overlap between any comparisons in the Venn diagrams. *represents genes that are not expressed in the blank treatment group.

Another observation is that some DEGs in XXF adipose and XXM liver are only expressed upon hormone treatment (estradiol and testosterone) but not when no hormone (blank) was administered (**Figure 3A; 3F**), suggesting their tight regulation by sex hormones. These include *Lcn13* (Lipocalin-13, involved in glucose metabolism) and *Slco1a1* (Solute Carrier Organic Anion Transporter 1A1, important in Na^+^ independent transport of anions such as prostaglandin E2 and taurocholate) in XXM liver, and *Mpp5* (Membrane Palmitoylated Protein 5, important in cell junction organization and PI3K-AKT signaling) and *Rfwd3* (Ring finger and WD repeat domain 3, important in ligase activity and p53 binding) in XXF adipose.

#### Top DEGs specific for estradiol and testosterone for liver and adipose tissue

Besides the overlapping DEGs between hormones, we investigated the top DEGs that were specific for each hormone for each tissue (**Table 1, Table 2**). In the liver, *Elovl3* (Elongation of Very Long Chain Fatty Acid Elongase 3; not expressed in the blank or estradiol treatment group) and *Cux2* (Cut Like Homeobox 2, important in DNA binding; not expressed in the blank or estradiol treatment group) were among the top upregulated and downregulated DEGs affected by testosterone across the four genotypes, respectively. By contrast, no consistent DEGs were found across all genotypes in the estradiol treated group in liver tissue.

In adipose tissue, the top DEGs for testosterone and estradiol generally varied across genotypes. Limited consistency in testosterone DEGs was seen for *Mt2* (Metallothionein-2, important in the detoxification of heavy metals) between XXM and XYM, and for estradiol treatment, consistency was seen between XXM and XYM for *Hpca* (Hippocalcin, involved in calcium binding) and between XXF and XYM for *Thbs1* (Thrombospondin 1, important in platelet aggregation).

### Identification of potential regulators of sex-biasing factors

#### Transcription factor (TF) network analysis

To understand the regulatory cascades that explain the large numbers of sex-biased genes affected by hormone treatments (**Figure 3**), we performed TF analysis using as input DEGs that passed an FDR<0.05 from 1WA specific to testosterone effects in the liver and estradiol effects in adipose tissue (**Table 1; Table 2**). For the testosterone liver DEGs, we identified 67, 66, 60 and 62 TFs for XYM, XXM, XXF and XYF respectively (**Figure 4A-D; Supplementary Table S6**). As expected, we captured gonadal hormone receptors including Androgen Receptor (*AR*) as a highly ranked TF in all genotypes and estrogen receptors (*ESR1, ESR2, ESRRA)* to be TFs with lower rank. We also found *NR3C1* (Nuclear receptor subfamily 3; the glucocorticoid receptor important for inflammatory responses and cellular proliferation) to be among the top 5 TFs for all four genotypes and the top-ranked TF for XXF and XYF, which is consistent with a female bias for this TF found in the GTEx study (Oliva et al. 2020). A number of circadian rhythm TFs were found throughout all genotypes in the liver including *CRY1*, *CRY2*, *PER1*, and *PER2*, which is consistent with sex differences in body clocks (Anderson and FitzGerald 2020). Additional consistent TFs for testosterone effect in liver across multiple genotypes, where sex bias has been documented to some extent previously, include *FOXA1/2*, *XBP1*, *HNF4A*, *SPI1*, and *CTCF*.

**Figure 4.**
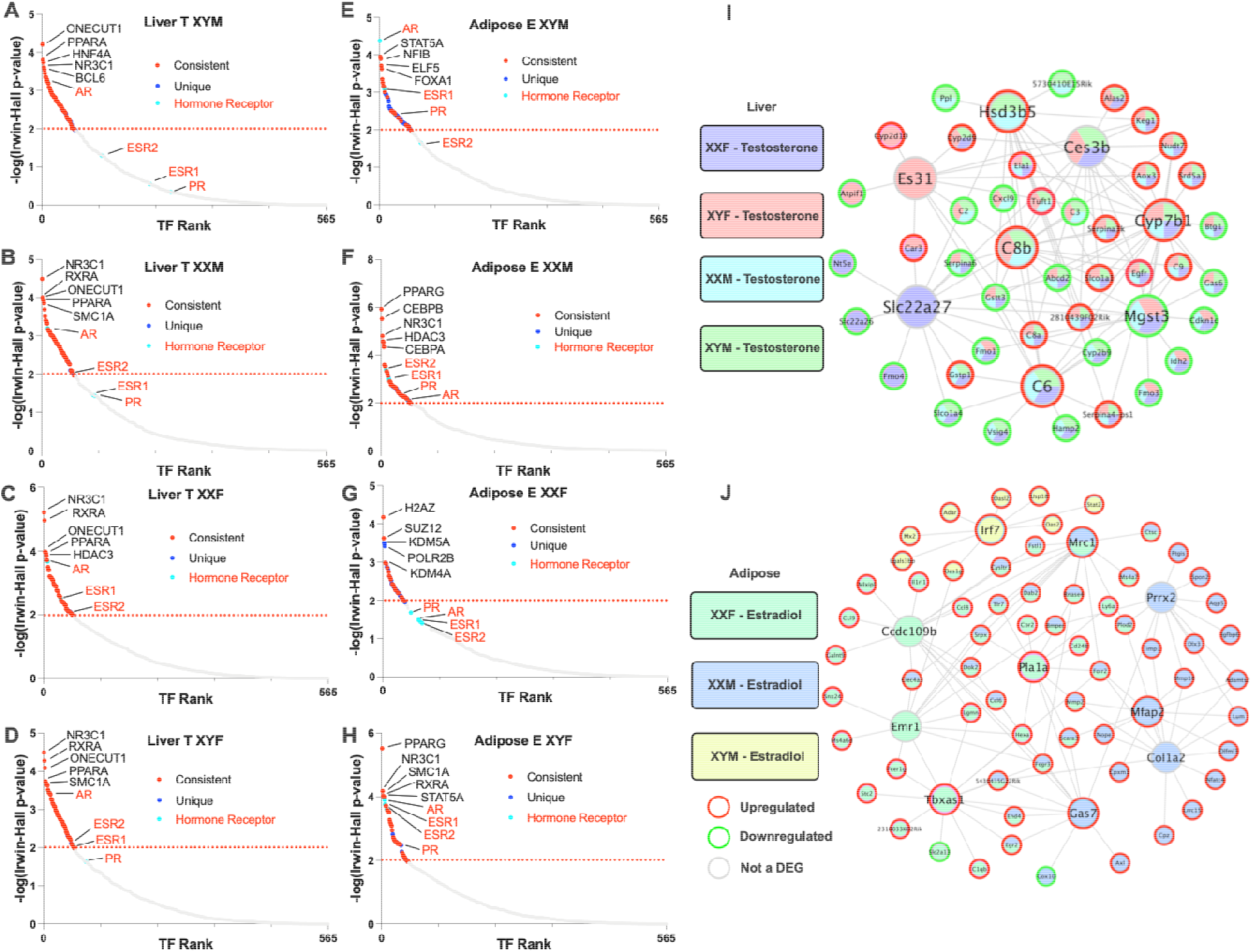
Transcription factor analysis (A-H) and key driver analysis (I-J) of DEGs informed by estradiol and testosterone treatment in liver and adipose. **A-D** represents TF analysis for liver. **E-H** represents TF analysis for adipose. For the TF network we utilized DEGs (FDR <0.05) from our 1WA for testosterone and estradiol treatment analysis using the BART tool, where a TF was considered significant by an Irwin-Hall p<0.01 analogous to −log10(p-value) = 2. Red color signifies the TF is present in at least one other genotype and the blue color signifies if the TF is only present in the given genotype. Turquoise color and red font denote a hormonal receptor relevant to testosterone and estradiol. Labelled TFs showcase the Top 5 by rank and additional hormonal receptors. **I.** Liver gene regulatory network (GRN). **J.** Adipose GRN. For GRN construction we overlaid DEGs (FDR <0.05) from our post hoc 1WA for testosterone and estradiol treatment onto our previously built adipose and liver Bayesian networks utilizing a KDA analysis from the Mergeomics package. We visualized the top 5 KDs for the testosterone or estradiol DEGs from each genotype group. KDs are labeled as larger nodes and DEGs as smaller nodes. Direction of DEGs is annotated with red or green borders for upregulation or downregulation, respectively.

An analysis of TFs that may mediate estradiol effects in adipose tissue identified 64, 61, 44 and 53 TFs for XYM, XXM, XXF and XYF respectively (**Figure 4E-H; Supplemental Table S7**). We found *ESR1* and *ESR2* as consistent TFs throughout the genotypes, except for XYF, where no classical estradiol or androgen receptor TF was captured. We also identified *AR* as a top TF in XYM and XYF. Notably, we found many TFs across our genotypes to be consistent with the TFs for female-biased genes in the Anderson et al. human adipose study (Anderson et al. 2020). Out of their top 20 ranked TFs for female-biased genes, we found 17 in our results for estradiol treatment in our genotypes, including *ESR1*, *H2AZ*, *SUZ12*, *KDM2B*, *CEBPB* and *PPARG*. When comparing TFs for consistency across genotypes, the top TFs were generally consistent, except *KDM5A*, *POLR2B*, *KMT2C* and *CLOCK* were particular to XXF.

When looking into the TFs that mediate estradiol’s effects in XYM for potential male-biased regulation in adipose tissue, we found matches with 13 of the top 20 TFs from the Anderson et al. human adipose study, with top TFs including *AR*, *CTCF*, *SMC1A*, *EZH2*, *ESR1*, *RAD21* and *TP63*, many of which were also consistent with additional studies in both mouse (Matthews and Waxman 2019; Anderson et al. 2020) and two studies of humans including the GTEx study (Anderson et al. 2020; Oliva et al. 2020).

#### Gene regulatory network analysis

An alternative and complementary approach to the TF analysis above is to utilize a gene regulatory network approach to decipher the key drivers (KDs) that may drive sex-biased gene alterations in each genotype based on the DEGs found in 1WA (**Table 1; Table 2**). We note that these KDs did not overlap with the TFs identified above due to the incorporation of genetic regulatory information in network construction.

In the liver (**Figure 4I**), we saw a large overlap between the KDs predicted across all four genotypes for testosterone treatment, but no KDs captured for estradiol treatment. We identified *Cyp7b1*, which is important in converting cholesterol to bile acids and metabolism of steroid hormones, as among the top 5 KDs for all genotypes. *Mgst3* (involved in inflammation), *C6* and *C8b* (complement genes), and *Ces3b* (xenobiotics detoxification) were top 5 KDs for 3 of the 4 genotypes (**Figure 4I**). We also identified KDs specific to particular genotypes (**Supplemental Table S8**) such as *Es31* (xenobiotics detoxification) for female gonads, *Slc22a27* (anion transport) for XXF, *Serpina6* (inflammation) for XYF, and *Hsd3b5* (steroid metabolism) for male gonads. Among these KDs, *Slc22a27* was previously found to be expressed predominantly in females and *Hsd3b5* and *Cyp7b1* were male specific (Adams et al. 2015), thus agreeing with our results.

For estradiol, 31 KDs were found for adipose tissue DEGs from the XXF, XXM and XYM genotypes (**Figure 4J; Supplemental Table S9**). The KDs included *Mrc1* (response to infection), which is the only overlapping top KD between genotypes XXF and XXM. KDs that were more highly ranked for XXM but still statistically significant in XXF included genes involved in extracellular matrix organization (*Prxx2*, *Mfap2*, *Col1a2*, and *Gas7*), and those specific to XXF are relevant to lipid synthesis/metabolism (*Tbxas1*, *Pla1a*) and immune function (*Emr1* and *Ccdc109b*). *Irf7* is the only KD for XYM, which has been recently suggested to be a TF in adipocytes with roles in adipose tissue immunity as well as obesity (Kuroda et al. 2020).

### Disease association of the genes affected by sex-biasing factors

Finally, to test the disease relevance of the genes affected by sex-biasing factors, we used a marker set enrichment analysis (MSEA; **details in Methods/Supplemental Methods**) to detect whether the DEGs highlighted in the 1WA overlap with genes previously identified to have SNPs associated with diseases/pathogenic traits by GWAS. In brief, we take the full summary statistics for each of the GWAS datasets, and then map each of the GWAS SNPs to genes using liver and adipose eQTLs. These mapped genes represent disease-associated genes informed by GWAS. Using the mouse orthologs of these human GWAS disease genes, we then look for enrichment of sex-biased DEGs from FCG to connect the genes affected by individual sex factors with human disease genes. Of the 73 disease/traits screened for which full GWAS summary statistics is available (**Supplemental Table S10)**, we focused on two broad categories, “cardiometabolic” (**Figure 5A;5B**) and “autoimmune” (**Figure 5C;5D**), both of which are known to show sex differences. The cardiometabolic category included Coronary Artery Disease (CAD), Type 2 Diabetes (T2D), fasting glucose level, BMI in women, BMI during childhood, BMI, total cholesterol (TC) level, triglyceride (TG) level, low-density lipoprotein (LDL) cholesterol level, and high-density lipoprotein (HDL) cholesterol level **(Figure 5A-B)**. The autoimmune category included Irritable Bowel Disease (IBD), Ulcerative Colitis (UC), Crohn’s Disease (CD), and Type 1 Diabetes (T1D) (**Figure 5C; 5D**). For hormone DEGs, we focused on those that are directly relevant to the general human population to understand how testosterone or estradiol can affect disease outcomes on XYM (physiological males) or XXF (physiological females).

**Figure 5.**
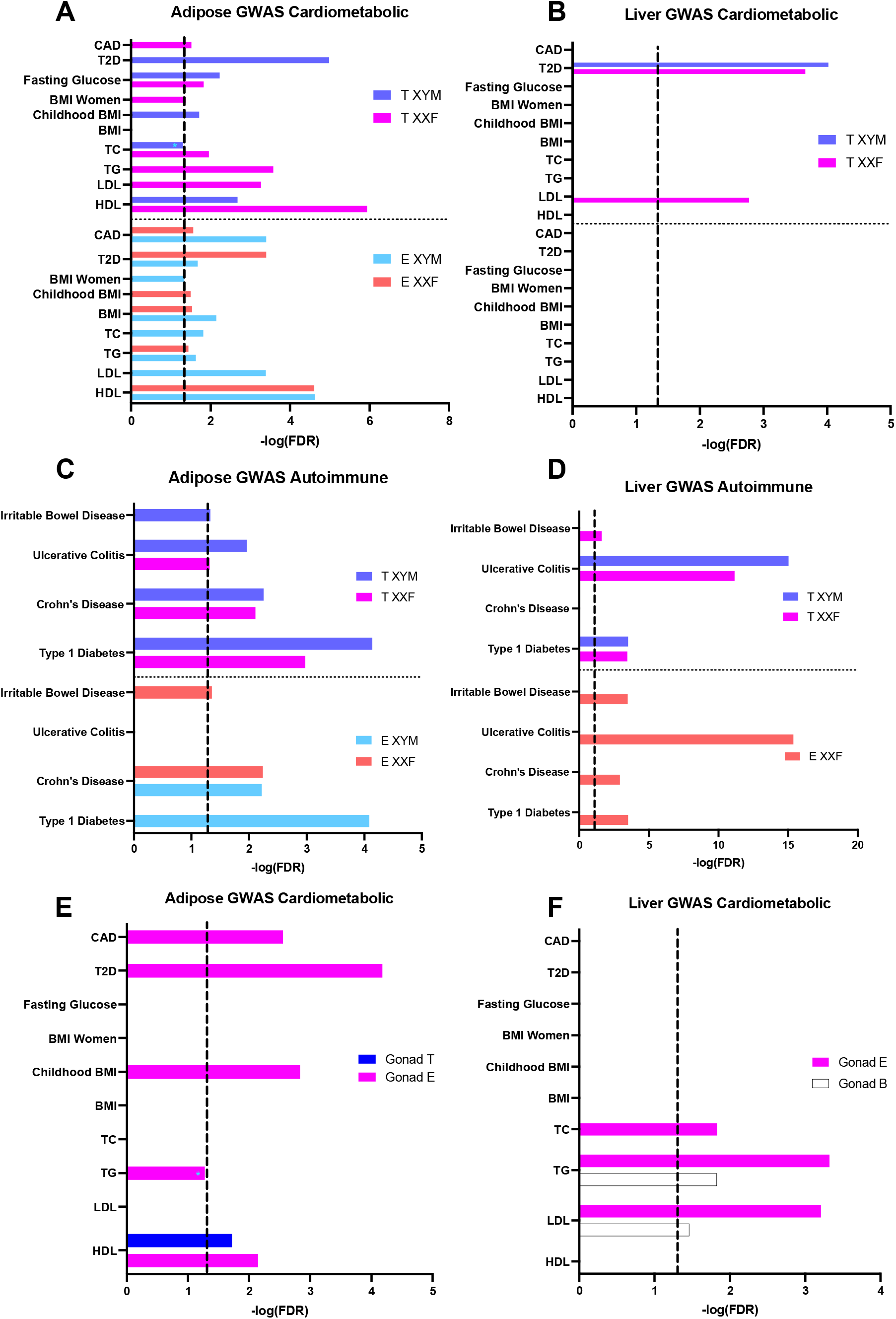
Bar graphs showing enrichment of the hormone DEGs and gonadal DEGs for known cardiometabolic and autoimmune diseases based on MSEA analysis. **A.** Bar graph for association of adipose DEGs from the post hoc one-way ANOVA for ten cardiometabolic diseases/traits. **B.** Bar graph for association of liver DEGs from the post hoc one-way ANOVA with cardiometabolic diseases/traits. **C.** Bar graph for association of adipose DEGs from the post hoc one-way ANOVA for five autoimmune diseases. **D.** Bar graph for association of liver DEGs from the post hoc one-way ANOVA for five autoimmune diseases. **E.** Bar graph for association of adipose DEGs from the two-way ANOVA for ten cardiometabolic diseases/traits. **F.** Bar graph for association of liver DEGs from the two-way ANOVA f ten cardiometabolic diseases/traits. **A-D**. DEGs at an FDR <0.05 derived from the posthoc one-way ANOVA were tested against genetic association signals with cardiometabolic and autoimmune diseases and traits. **E** and **F** DEGs at an FDR < 0.05 were tested against genetic association signals with cardiometabolic diseases. Dotted line signifies FDR <0.05 and * denotes enrichment minimally below the FDR< 0.05 cutoff.

#### Disease association for hormone DEGs

When adipose tissue and cardiometabolic diseases were considered **(Figure 5A; Supplemental Table S11)**, testosterone DEGs from XYM were enriched for GWAS signals for 5 cardiometabolic diseases (T2D and childhood BMI being unique), and those from XXF were associated with 7 cardiometabolic diseases (CAD, BMI in women, TG, and LDL being unique), with 3 shared (Fasting glucose, TC and HDL) between genotypes. In terms of autoimmune diseases (**Figure 5C; Supplemental Table S12**), testosterone DEGs in XYM were enriched for all diseases and those in XXF for all but IBD. For estradiol DEGs, we found 8 cardiometabolic diseases enriched in XYM (BMI in women, TC, and LDL being unique) and 6 cardiometabolic diseases enriched in XXF (childhood BMI being unique), with 5 shared (CAD, T2D, BMI, TG and HDL) between genotypes. For autoimmune diseases, estradiol DEGs in XYM are enriched for CD and T1D, and those in XXF for CD and IBD. UC appears to be a unique association with testosterone DEGs whereas CD is associated with adipose DEGs of both hormones.

When liver tissue and cardiometabolic traits were analyzed (**Figure 5B; Supplemental Table S13**), we see low level enrichment but specificity of testosterone DEGs for both T2D and LDL, whereas no estradiol DEGs were enriched for any of the ten traits. For the association of liver DEGs with autoimmune disease (**Figure 5D; Supplemental Table S14**), we found that testosterone DEGs in XYM to be related to UC and T1D, whereas DEGs in XXF to be related to IBD, UC and T1D. Interestingly, estradiol DEGs from XYM had no association with autoimmune diseases but DEGs in XXF were associated with all autoimmune diseases. Taken together, liver DEGs in XXF (gonadal females) affected by both testosterone and estradiol were enriched for autoimmune diseases, implying the modifiable effect of either hormone in these diseases in females.

Overall, the disease association analyses identified adipose DEG sets altered by both hormones for both cardiometabolic and autoimmune processes. For liver DEGs, the most significant associations were with autoimmune diseases and subtle T2D and LDL associations were identified for liver testosterone DEGs. It is unclear, however, whether the associations would be pathogenic or protective.

#### Disease association for gonadal sex DEGs

We also used MSEA to detect whether gonadal DEGs highlighted in 2WA (FDR<0.05) overlap with genes previously identified to have SNPs associated with diseases/pathogenic traits by GWAS (**Figure 5E-F**). For adipose tissue, the gonadal DEGs were particularly enriched in cardiometabolic disease/trait GWAS when on an estradiol background, capturing significance in CAD, T2D, Childhood BMI and HDL traits. Gonadal DEGs on a testosterone background only showed association with HDL and no disease association was captured when no hormone (blank) is the background (**Supplemental Table S15**). In regard to the liver, gonadal DEGs on an estradiol background were enriched for TC, TG, and LDL GWAS signals (**Supplemental Table S16**).

#### Disease association for sex chromosome DEGs

Due to the low number of DEGs captured for the sex chromosome effect, no enrichment results are possible through MSEA, therefore we queried whether sex chromosome DEGs have been previously implicated in human diseases by overlapping the DEGs (FDR<0.05 from 2WA) with candidate genes from the GWAS catalog for 2203 traits. Both adipose tissue and liver DEGs demonstrating sex chromosome effects overlapped with GWAS candidates for both cardiometabolic and autoimmune disease (**Supplemental Table S17**) as well as various other diseases (**Supplemental Table S16**).

#### Disease association for DEGs showing interactions among sex-biasing factors

Similarly, overlapping DEGs that showed interactions among the sex-biasing factors (**Supplemental Table S2, S4**) with GWAS candidate genes, demonstrated that both liver and adipose DEGs affected by interaction between gonad and hormone, and between sex chromosome and gonad overlapped with numerous autoimmune and cardiometabolic traits and disease (**Supplemental Table S19**).

## DISCUSSION

The variation in physiology and pathophysiology between sexes is established via the modulatory effects of three main classes of sex-biasing agents: sex chromosome complement (XX vs. XY), long-lasting organizational effects of hormones secreted by the gonads at critical periods such as prenatally or at puberty, and the acute, transient activational effects of circulating hormones that wax and wane at various life stages (Arnold 2009). The manifestations of these sex-dependent modulators impact disease incidence and severity, including metabolism-related diseases and autoimmune diseases (Voskuhl and Gold 2012; Link and Reue 2017). In this study, we separated the effects of these sex-biasing components using the FCG model, thus enabling the analysis of each contributing factor as well as their interactions in altering gene expression in inguinal adipose and liver tissues, which are relevant in systems metabolism and immunity (**Figure 1**).

Our data revealed distinct patterns between tissues in the relative contribution of each sex-biasing factor to gene regulation (**Table 1; Table 2**). In particular, the liver transcriptome is mainly affected by acute effects of testosterone, followed by acute effects of estradiol, organizational effect of gonadal sex, and sex chromosome complement, whereas inguinal adipose gene expression is primarily regulated by acute effects of estradiol, followed by gonadal sex, acute effects of testosterone, and sex chromosome complement. The genes and pathways regulated by the sex-biasing factors are largely different between factors, although metabolic, developmental, and immune functions can be regulated by both activational effects of sex hormones and gonadal sex (organizational effects). Sex chromosome effects were primarily associated with genes that reside on X and Y chromosomes, along with a handful of autosomal genes involved in inflammation and metabolic processes that are downstream of the sex-biasing effects of X and Y genes. Cell deconvolution analysis supports that sex-biasing factors influence the proportion of diverse cell populations such as immune cells, hepatocytes, and dividing cells, suggesting that cellular composition changes may partially explain the observed genes and pathways. Lastly, the liver and adipose tissue genes affected by the sex-biasing factors were found to be downstream targets of numerous TFs and network regulators, not just the sex hormone receptors, and show association with human cardiometabolic and autoimmune diseases (**Figure 6**).

**Figure 6.**
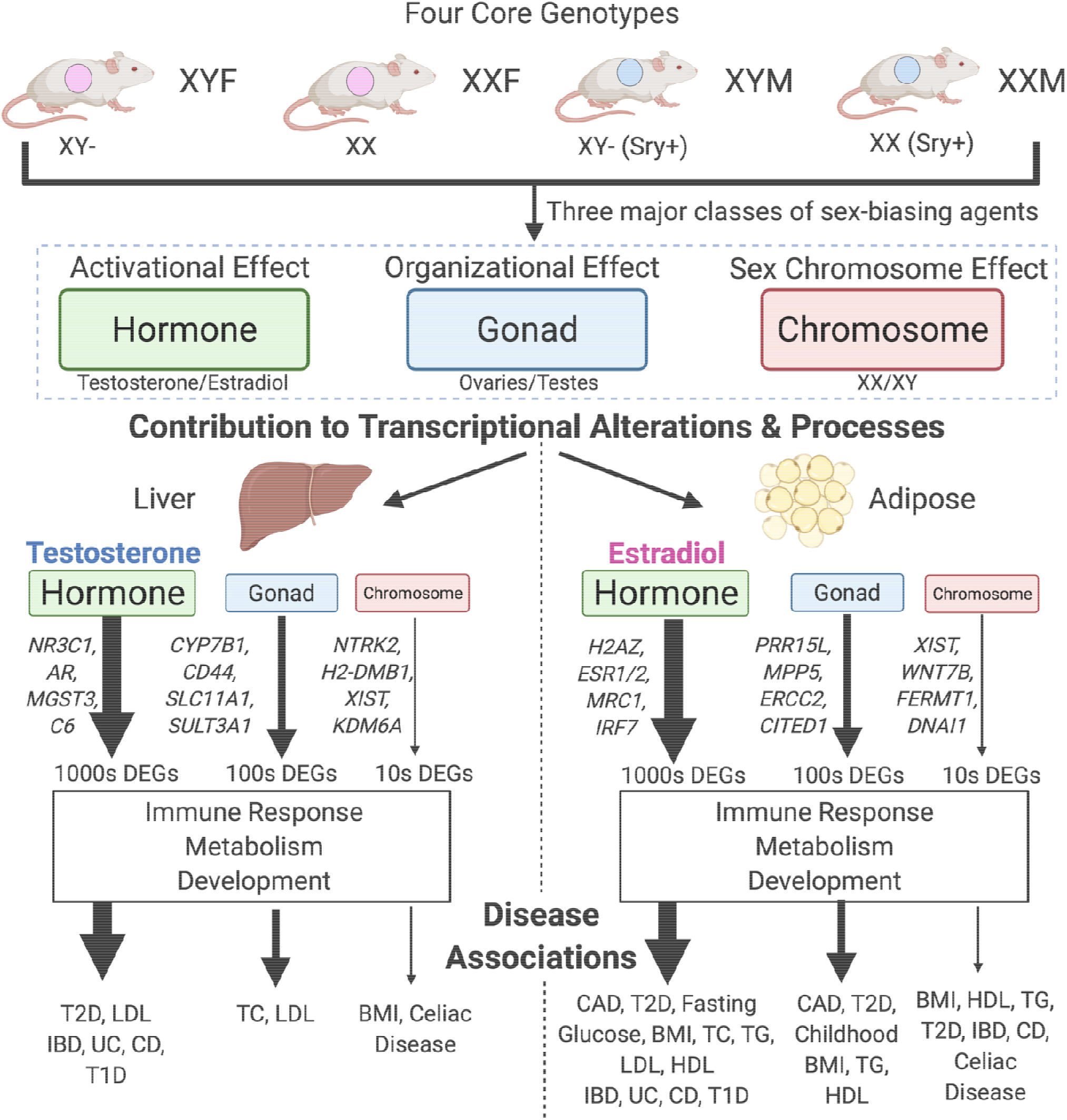
Study Summary. Utilizing the FCG model, we separated the effects of three major classes of sexbiasing agents and uncovered their relative contribution to transcriptional alterations in the liver and adipose tissue, the resulting biological processes enriched and finally the diseases associated.

Previously, sex differences in the liver transcriptome have been largely attributed to sex differences in the circadian rhythm and levels of Growth Hormone, which are established because of perinatal organizational masculinization of hypothalamo-pituitary mechanisms controlling Growth Hormone (Mode and Gustafsson 2006; Waxman and Holloway 2009; Sugathan and Waxman 2013). Genes regulated in this manner would be expected to appear in the gonadal effect DEGs. Our results suggest, however, that the acute activational effects of gonadal hormones might be a more important influence, because of the larger number of testosterone or estradiol DEGs compared to gonad DEGs. Our results support previous evidence that removal of gonadal hormones in adulthood eliminates most sex differences in mouse liver gene expression (van Nas et al. 2009; Norheim et al. 2019), and that liver-specific knockout of estrogen receptor alpha or androgen receptor altered genes that underlie sex differences in the liver transcriptome (Zheng et al. 2018). It is possible that the effects of gonadal steroids during adulthood are required for some of the organizational effects of testosterone mediated via Growth Hormone action. In contrast to liver, gonadectomy does not eliminate sex differences in the adipose transcriptome (Norheim et al. 2019), which agrees with our finding that the organizational effects of gonads play a strong role, in addition to estradiol, in adipose gene regulation.

Although we observed a large effect of hormonal treatment relative to gonadal sex and sex chromosome complement for both tissues (**Figure 2A-F**), the striking tissue-specificity for each of the sex-biasing factors observed here highlights that individual tissues have unique sex-biased regulatory mechanisms. Circulating levels of testosterone play a major role in determining sex differences in liver gene expression patterns (**Figure 2A**), confirming earlier studies (van Nas et al. 2009; Norheim et al. 2019), whereas levels of estradiol as well as gonadal sex are the main factors to drive sex differences in adipose transcriptome (**Figure 2D**). For adipose tissue, the stronger estradiol effect than testosterone is consistent with evidence that ERs are expressed at higher levels than ARs in adipose tissue, as well as having higher overall expression in females (Shen and Shi 2015b). In the liver, ERs are equally distributed between males and females but are generally expressed at a low level (Eisenfeld and Aten 1987). However, ARs are present at much higher concentrations (Eisenfeld and Aten 1987) and higher expression levels in males compared to females (Shen and Shi 2015b); this agrees with our TF analysis ranking ARs more highly than ERs in the liver for all genotypes.

We found that the gonadal sex factor primarily affects developmental pathways, cell adhesion, and metabolic pathways in adipose (**Table 2; Figure 2H**), which corroborates past evidence indicating that early gonadal sex status and associated hormonal release play critical roles in the development of sex differences and disease outcomes (Leung et al. 2004; Varlamov et al. 2012; Shen and Shi 2015a).

Compared to the organizational gonadal sex effects and activational hormone effects, the sex chromosome effects were minimal, and no coherent pathways were found for the sex chromosome-driving DEGs (**Table 1; Table 2**) or co-expression modules (**Figure 2G-H)**. The DEGs include those known to escape X inactivation (*Kdm6a*, *Eif2s3x*, *Ddx3x*) (Chen et al. 2012; Berletch et al. 2015) and their Y paralogues (*Eif2s3y*, *Ddx3y*). The X escapees are expressed higher in XX than XY cells, causing sex differences in several mouse models of metabolic, immune, and neurological diseases (Kaneko and Li 2018; Itoh et al. 2019; Davis et al. 2020; Link et al. 2020). Comparison between XXF and XYF revealed that immune and arrhythmogenic right ventricular cardiomyopathy (ARVC) related pathways are enriched among the DEGs affected by sex chromosomes when gonadal hormonal levels are absent (blank group) in these mice that were never masculinized by organizational effects of testicular androgens. This may be relevant to the postmenopausal state in women, which is characterized by an increase in risk of heart and autoimmune disease, potentially driven by sex chromosomal effects.

As our comparative analysis of the three classes of sex-biasing factors clearly determined that the activational effects of gonadal hormones are the dominant factors, we further investigated potential upstream regulatory factors that may control the sex-biased genes, using a gene regulatory network analysis and a TF analysis, revealing both expected and novel findings. In concordance with the importance of hormonal effects and consistent with recent human studies including GTEx searching for tissue-specific sex bias (Anderson et al. 2020; Oliva et al. 2020), TFs for hormone receptors (*AR* and *ESR1/2*) were captured in the majority of genotypes (**Figure 4A–4H**). Beyond the major hormonal receptors, within the liver numerous circadian related TFs were captured (*PER1*, *PER2*, *CRY1* and *CRY2*). Although it is known that males and females have differing biological clocks (Anderson and FitzGerald 2020), the contribution of hormones particularly in this rhythm is far from fully elucidated and our findings support that hormones need to be taken into account in liver circadian rhythm studies. In adipose tissue for estradiol treatment, however, we found that surprisingly the XXF genotype has no significant signal for ERs, which may imply that estradiol’s major contribution in adipose gene regulation is more importantly through TFs such as *H2AZ*, which have been shown to be essential for estrogen signaling and downstream gene expression (Gévry et al. 2009). In addition to TFs, we utilized a GRN analysis, revealing non-TF regulators. For the liver GRN (**Figure 4I**), key driver genes for testosterone DEGs are involved in immune processes (*Mgst3*, *C6*, *C8b*), steroid metabolism (*Cyp7b1* and *Hsd3b5*), and xenobiotic detoxification (*Ces3b* and *Es31*). In adipose tissue (**Figure 4J**), the network is dominated for estradiol signals likely due to the larger number of DEGs as compared to testosterone treatment. Here, we see far fewer shared key drivers across genotypes relative to the results in the liver with testosterone treatment, indicating that estradiol has more finely tuned interactions with the gonadal sex and sex chromosome genotypes than the broad effect of testosterone. The role of the numerous liver and adipose regulators uncovered in our analysis warrant further investigation.

Lastly, to provide context to the health relevance of the liver and adipose sex-biasing DEG sets, we looked for GWAS association of these genes with human diseases/traits. We found that hormone-affected genes in adipose tissue were enriched for genetic variants associated with numerous cardiometabolic diseases/traits, but the enrichment was weaker for the liver DEGs (**Figure 5A; 5B**). CAD and its related traits (HDL, LDL, TG, TC, BMI and fasting glucose) showed significant and widespread enrichment in adipose DEGs for both testosterone and estradiol. Another important area of sex difference is found within autoimmunity, which occurs more in females (Mauvais-Jarvis et al. 2020). While both adipose and liver DEGs from multiple hormone-genotype combinations were enriched for autoimmune diseases, the liver DEGs, particularly those from the XXF genotype, had more prominent autoimmune association. Beyond the hormonal DEG enrichment in human disease/trait, we also found that DEGs caused by gonad type from both adipose and liver are involved in cardiometabolic disease (**Figure 5E; 5F**). Finally, despite minimal DEGs captured for the sex chromosome effect as well as the interactions between each sex biasing factors, we found overlap of these DEGs with various disease traits. The DEGs underlying disease associations may explain the differential susceptibility of males and females to these major diseases, and warrant further investigation to distinguish risk versus protection through the genes identified in this study.

The analyses presented in this study show an extensive dissection of the relative contribution of three classes of sex biasing factors on liver and adipose gene expression, their associated biological processes and regulators, and their potential contribution to disease. Importantly, many of the genes identified in our study were replicated in independent human studies such as GTEx, and our mouse study offers unique insights into the particular sex-biasing factors (hormones, sex chromosomes, or gonads) that likely contribute to the sex-biased gene expression in humans. Despite retrieving numerous new insights, we acknowledge the following limitations. First, gonadectomy and subsequent treatment of hormones may have caused activational effects that do not match the effects of endogenous physiological changes in the same hormones, leading to more predominant activational effects being observed. Second, the relative effects of testosterone and estradiol are affected by the doses of each hormone used. Extensive further investigations, using numerous doses of each hormone, are required for detailed comparison of effects of the two hormones. Third, we used DEG counts as a measure of overall effect size to compare the various sex-biasing factors, which may be influenced by sample size and statistical power. Therefore, caution is needed when interpreting the results. However, when we fit the 3WA, ~60 samples/tissue were taken into consideration, with sample sizes of n=~20 per hormone (blank vs. T; blank vs. E), ~30 per sex chromosome (XX vs. XY), and ~30 per gonad (M vs. F). The sample sizes are comparable across sex-biasing factors and are adequate for mouse transcriptome studies with sufficient statistical power (Pawitan et al. 2005; Lin et al. 2010). Fourth, the comparison of mice with testes vs. ovaries does not map perfectly onto mice that had organizational effects of testicular vs. ovarian secretions because of the potential effects of the *Sry* transgene, which was present in tissues only of mice with testes. Lastly, only liver and inguinal adipose tissues were investigated, other tissues warrant examination in future studies.

Overall, our data revealed tissue-specific differential gene expression resulting from the three sex-biasing factors, thereby distinguishing their relative contributions to the differential expression of key genes in a variety of clinically significant pathways including metabolism, immune activity, and development. Importantly, in addition to establishing the critical influence of hormones and their effect on the transcriptome in a tissue specific manner, we also uncovered and highlighted the underappreciated role of the sex chromosomal effect and organizational gonadal effect as well as interactions among sex-biasing factors in global gene regulation. Our findings offer a comprehensive understanding of the origins of sex differences, and each of their potential associations with health and disease.

## METHODS

### Overall Study Design

In this study we used FCG mice on a C57BL/6J B6 background (B6.Cg-TgSry2Ei Srydl1Rlb/ArnoJ, Jackson Laboratories stock 10905; backcross generation greater than 20), bred at UCLA (De Vries et al. 2002; Burgoyne and Arnold 2016). Gonadal females and males were housed in separate cages and maintained at 23°C with a 12:12 light: dark cycle. Animal studies were performed under approval of the UCLA Institutional Animal Care and Use Committee.

A total of 60 FCG mice, representing 4 genotypes (XXM, XXF, XYM, XYF), were gonadectomized (GDX) at 75 days of age and implanted immediately with medical grade Silastic capsules containing Silastic adhesive only (blank control, (B) or testosterone (T) or estradiol (E) (**details in Supplemental Methods**). Mice were euthanized 3 weeks later; liver and inguinal adipose tissues were dissected, snap frozen in liquid nitrogen and stored at −80°C for RNA extraction and Illumina microarray analysis.

### Genotyping

DNA was extracted from tails or ears using Chelex resin (Bio-Rad, Hercules, CA). The genotype of mice was determined by PCR using the primers described in (Itoh et al. 2015) and (Burgoyne and Arnold 2016).

### RNA isolation, microarray hybridization, and quality control

RNA from liver and inguinal adipose tissue was isolated using Trizol (Invitrogen, Carlsbad, CA). Individual samples were hybridized to Illumina MouseRef-8 Expression BeadChips (Illumina, San Diego, CA) by Southern California Genotyping Consortium (SCGC) at UCLA. Two adipose samples were removed from the total of 60 after RNA quality test (degradation detected) and did not go for microarray. Principal Component Analysis (PCA) was used to identify three outliers among the adipose sample, which were removed from subsequent analyses. PCA was conducted using the prcomp R package with the correlation matrix. Sample size for liver tissue was = 5/genotype/treatment; for inguinal adipose the sample size is XXM = 14 (B = 5, E = 4, T = 5), XXF = 12 (B = 3, E = 5, T = 4), XYM = 14 (B = 4, E = 5, T = 5), and XYF = 15 (B = 5, E = 5, T = 5).

### Identification of differentially expressed genes (DEGs) affected by individual sex-biasing factors

To identify DEGs, we conducted 3-way ANOVA (3WA), 2-way ANOVA (2WA), and 1-way ANOVA (1WA) using the aov R function. The 3WA tested the general effects of 3 factors of sex chromosomes, gonad, and hormonal treatments, as well as their interactions, using the formula “gene expression ~ gonad + chromosome + hormone + gonad:chromosome + gonad: chromosome + chromosome:hormone + gonad:chromosome:hormone”. Two sets of 3WA were conducted, one comparing T vs. B, and the other E vs. B, to discriminate separate effects of the two hormones. The 2WA tested the effects of sex chromosomes and gonads as well as their interactions within each hormonal treatment group (T, E, or B) separately using the formula “gene expression ~ gonad + chromosome + gonad:chromosome”. For 1WA, we tested the effects of T (comparing T vs. B) and E (comparing E vs. B) within each genotype, using the formula “gene expression ~ treatment”. We used different sets of ANOVA analysis because each address *a priori* questions of this study, and offers different perspectives on the roles of the three factors. The different sets of ANOVAs offer different levels of granularity of sex differences from broad overviews to specific effects on a sex-biasing factor in a given setting. For example, the 3-way ANOVA provides a general view of the effects of hormones, gonads and chromosomes as well as the interactions, and rationalizes further analyses. To test for sex chromosome and gonadal effects within a specific hormone condition, we carried out three sets of 2-way ANOVAs in which hormone condition was fixed (e.g., either blank, or estradiol, or testosterone). To further capture the effects of hormones on each specific genotype, we used a 1-way ANOVA with post hoc analysis in each genotype. Multiple testing was corrected using the Benjamini-Hochberg (BH) method, and significance level was set to FDR <0.05 to define significant DEGs. Suggestive DEGs at FDR<0.1, p<0.01 and p<0.05 were also retrieved for analyses that are less sensitive to DEG cutoffs, such as pathway analysis.

### Co-expression network construction and identification of co-expression modules affected by individual sex-biasing factors

As co-expression networks can reveal unique biology that cannot be retrieved by DEG analysis (Huan et al. 2015; Cordero et al. 2019; Zhao et al. 2019), we used the Multiscale Embedded Gene Co-expression Network Analysis (MEGENA) (Song and Zhang 2015), a method similar to WGCNA (Langfelder and Horvath 2008), to recognize modules of co-expressed genes affected by the three different sex-biasing factors (details in Supplementary Methods). The unique strength of MEGENA compared to WGCNA is that it allows a gene to be in multiple modules and produces smaller and more functionally coherent modules. The influence of each sex-biasing factor on the resulting modules was assessed using the first principal component of each module to represent the expression of that module, followed by 3WA, 2WA, 1WA tests and FDR calculation as described under the DEG analysis section to identify differential modules (DMs) at FDR <0.05 that are influenced by the various sex-biasing factors.

### Annotation of the pathways over-represented in the DEGs and DMs

For each of the DEG sets and DMs that were significantly affected by any of the three sex-biasing factors, we conducted pathway enrichment analysis against Gene Ontology (GO) Biological Processes and KEGG pathways derived from MSigDB (with the mouse genome as our background) using Fisher’s exact test, followed by BH FDR estimation. Pathways that had >5 overlapping genes with the DEGs or DMs and FDR < 0.05 were used to annotate the functions of the DEGs or modules.

### Gene regulatory network analysis

To predict potential regulators of the sex-biased DEGs, we used the Key Driver Analysis (KDA) function of the Mergeomics pipeline (Shu et al. 2016) and liver and adipose Bayesian networks. In brief, the Bayesian networks were built from multiple large human and mouse transcriptome and genome datasets (Yang et al. 2006; Wang et al. 2007; Emilsson et al. 2008; Schadt et al. 2008; Tu et al. 2012). To identify the hub or key driver (KD) genes within these tissue-specific networks, the KDA uses a Chi-square like statistic to identify genes that are connected to a significantly larger number of DEGs than what would be expected by random chance (Supplemental Methods for details). KDs represent prioritized regulatory genes based on network topology. Bayesian network key driver genes were considered significant if they passed an FDR<0.05. The Bayesian networks of top key drivers were visualized using Cytoscape (Shannon et al. 2003).

### Transcription factor (TF) analysis

To predict TFs that may regulate the sex-biased DEGs sets, we used the Binding Analysis for Regulation of Transcription (BART) computational method (Wang et al. 2018). We followed the tool’s recommendation of a minimum of 100 DEGs as input and an Irwin-Hall p-value cut off (p < 0.01) for identify TFs.

### Maker Set Enrichment Analysis (MSEA) to connect sex biasing DEGs with human diseases or traits

To assess the potential role of the DEGs affected by each of the sex-biasing factors in human diseases, we collected the summary statistics of human GWAS for 73 diseases or traits that are publicly available via GWAS catalog (MacArthur et al. 2017). SNPs that have linkage disequilibrium of r^2^>0.5 were filtered to remove redundancies. To map GWAS SNPs to genes, we used GTEx Version 7 eQTL data for liver and adipose tissues (Lonsdale et al. 2013) to derive tissue-specific genes potentially regulated by the SNPs. We then use the MSEA function embedded in Mergeomics (Shu et al. 2016) which extracts the disease association p-values for the mapped SNPs from the summary statistics of each of the human GWAS datasets. The disease association p-values of the SNPs representing the DEGs were then compared with those of the SNPs mapped to random genes to assess whether the DEGs contain SNPs that show stronger disease associations than random genes for each disease using a Chi-square like statistic (**details in Supplemental Methods**).

### Deconvolution of bulk liver and inguinal adipose tissue

We downloaded single cell RNA-seq data for mouse liver from GEO (GSE166178) and mouse inguinal adipose from GEO (GSE133486) as our reference datasets, and utilized the deconvolution tool CIBERSORTx (Newman et al. 2019) for each genotype under each hormone treatment. We used the Impute Cell Fractions function and ran for 100 permutations. Cell proportion estimates were compared across groups using 1WA with posthoc analysis to identify cell types influenced by sex hormones. and t-tests for gonads, and sex chromosomes.

## DATA ACCESS

All raw and processed sequencing data generated in this study have been submitted to the NCBI Gene Expression Omnibus (GEO; https://www.ncbi.nlm.nih.gov/geo/) under accession number GSE176033. R code used in the analysis is accessible via GitHub (https://github.com/XiaYangLabOrg/FCG).

## ACKNOWLEDGEMENTS

AA was supported by NIH grants NS043196, HD076125, and HD100298. XY was supported by NIH grants DK104363, DK117850, and HD100298. MB is supported by the American Heart Association Predoctoral Fellowship (829009) and UCLA IBP Edith Hyde Fellowship. We thank Emilie Rissman for advice.

## AUTHOR CONTRIBUTIONS

MB formal analysis, visualization and writing of the manuscript. XC experimental work and writing, YI formal analysis, YZ formal analysis and writing, YH formal analysis, CM formal analysis, BS formal analysis and writing. AA study design, resources, supervision, writing, review and editing. XY analysis design, supervision, writing, review and editing. KR study design, resources, writing, review and editing.

