## Supplemental Figures for "Relative contributions of sex hormones, sex chromosomes, and gonads to sex differences in tissue gene regulation": Supplemental_Fig_S9.pdf

### Liver & Adipose 3 Way ANOVA DEG Comparison between Testosterone (T) and Estradiol (E)

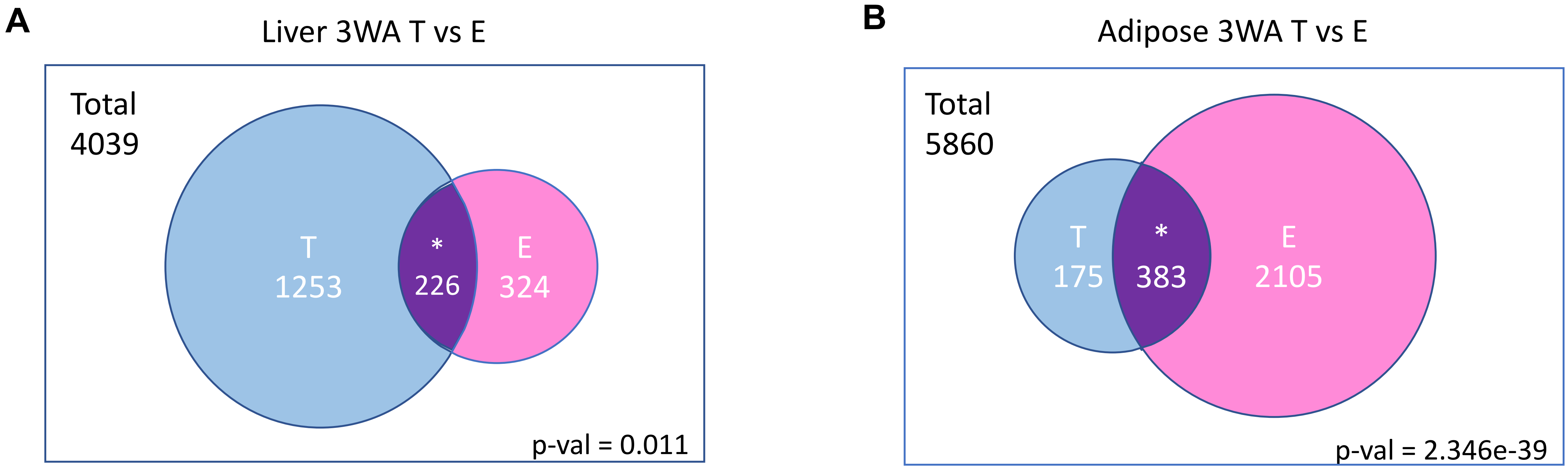


Top 5 genes overlapped: Sult3a1, C6, Pte2b, Serpina6, Serpina4-ps1

Top 5 genes overlapped: Gas6, Lrg1, Hsd11b1,Prtn3, Ces1f
