## Supplemental Figures for "Relative contributions of sex hormones, sex chromosomes, and gonads to sex differences in tissue gene regulation": Supplemental_Fig_S10.pdf

### Liver One-way ANOVA DEG Comparison between Testosterone (T) and Estradiol (E) in each genotype

Total # of genes: 307

#### Liver 1WA XXM

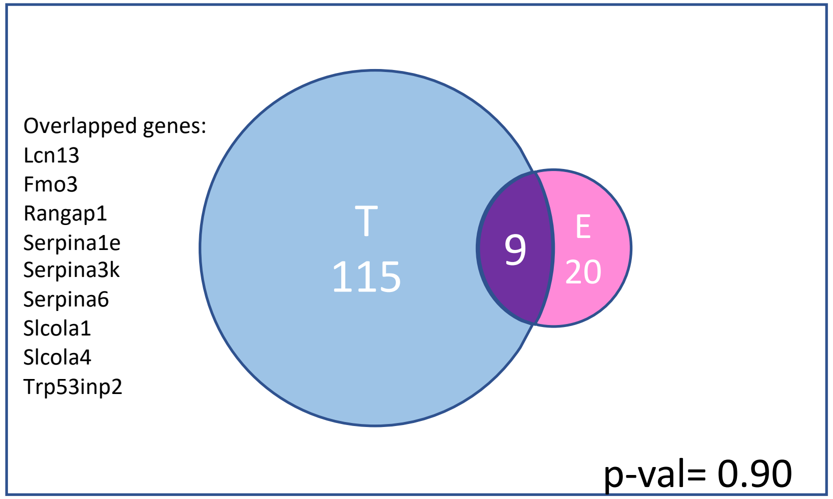


#### Liver 1WA XYM

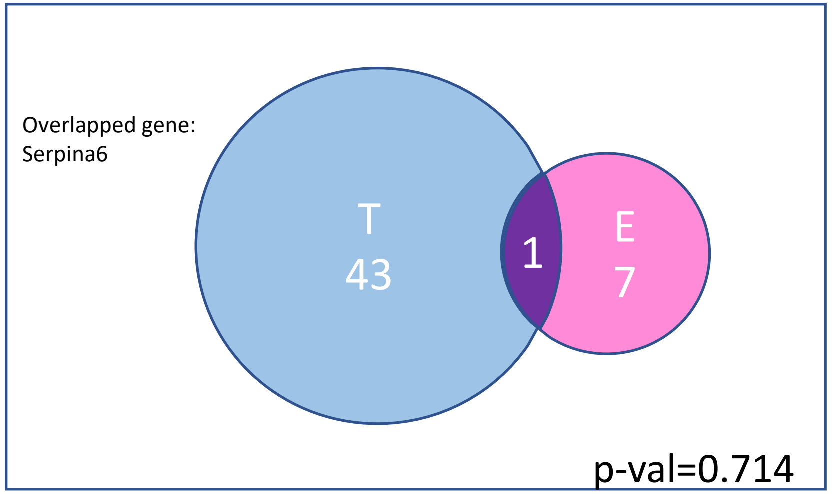


#### Liver 1WA XXF

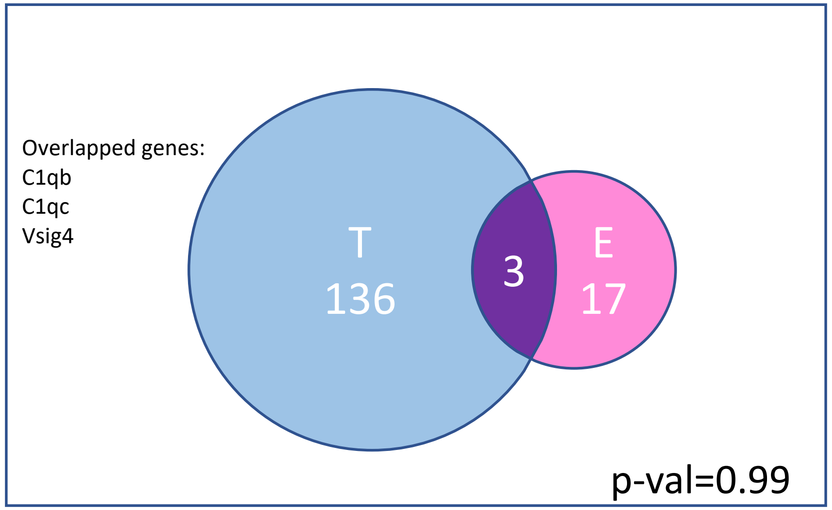


#### Liver 1WA XYF

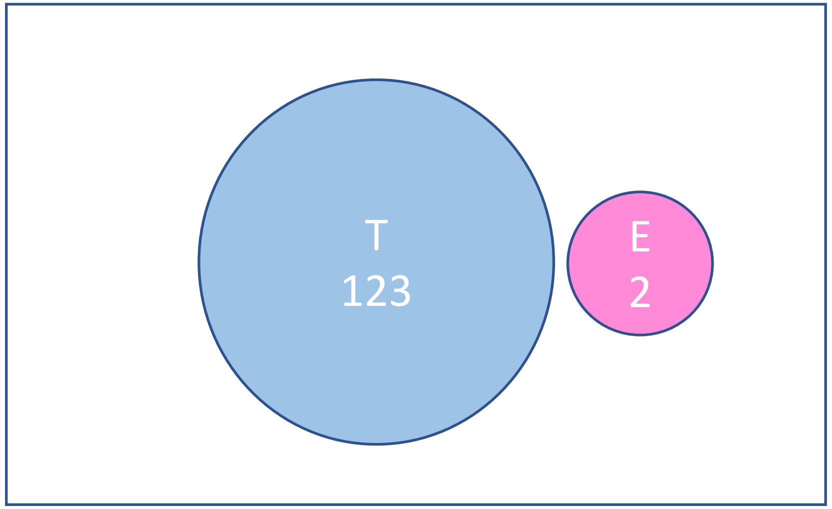

### Adipose One-way ANOVA DEG Comparison between Testosterone (T) and Estradiol (E) in each genotype

Total # of genes: 677

#### Adipose 1WA XXM

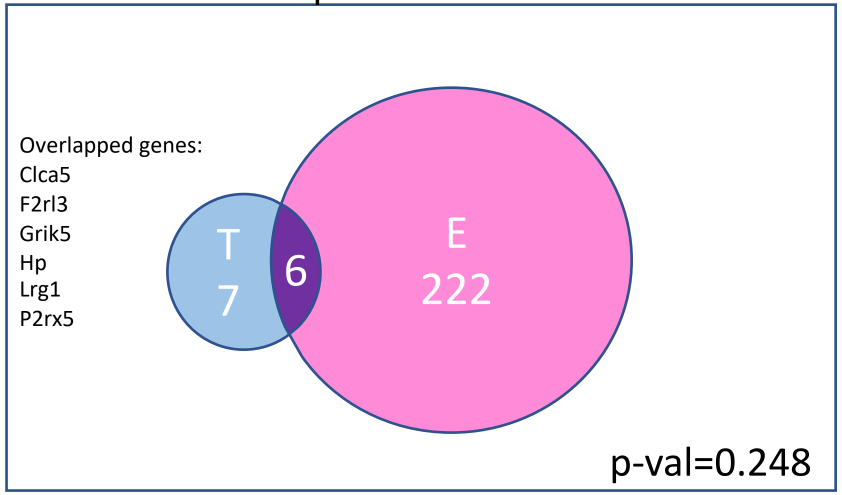


#### Adipose 1WA XYM

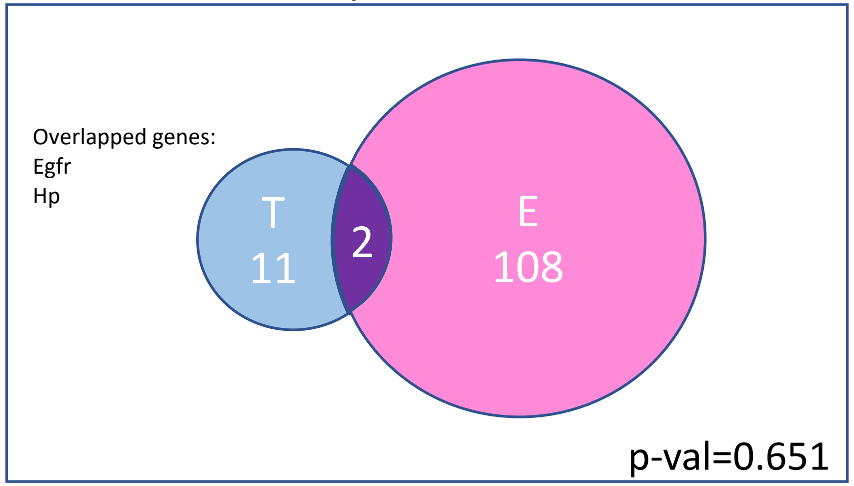


#### Adipose 1WA XXF

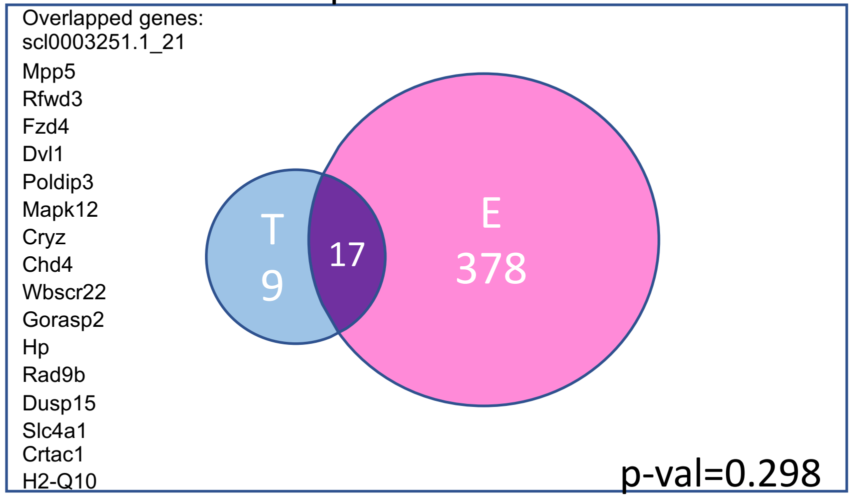


#### Adipose 1WA XYF

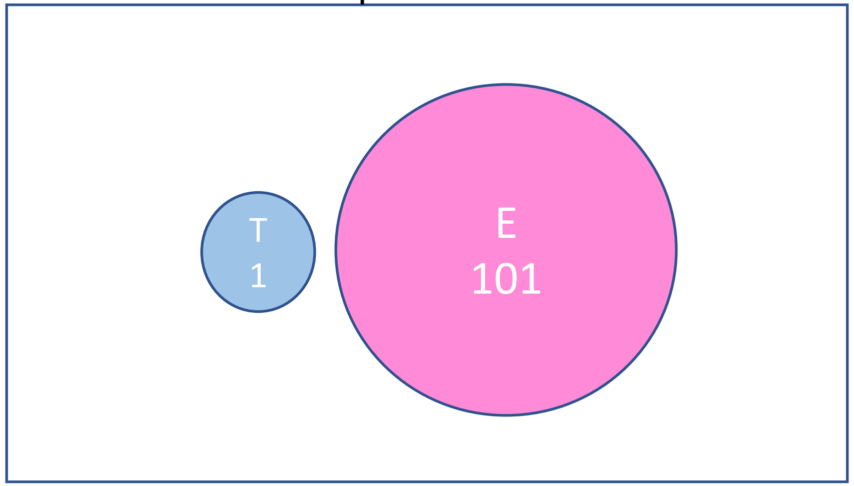
