## Supplementary figures and images for "Relative contributions of sex hormones, sex chromosomes, and gonads to sex differences in tissue gene regulation"

### Supplemental_Fig_S1.png

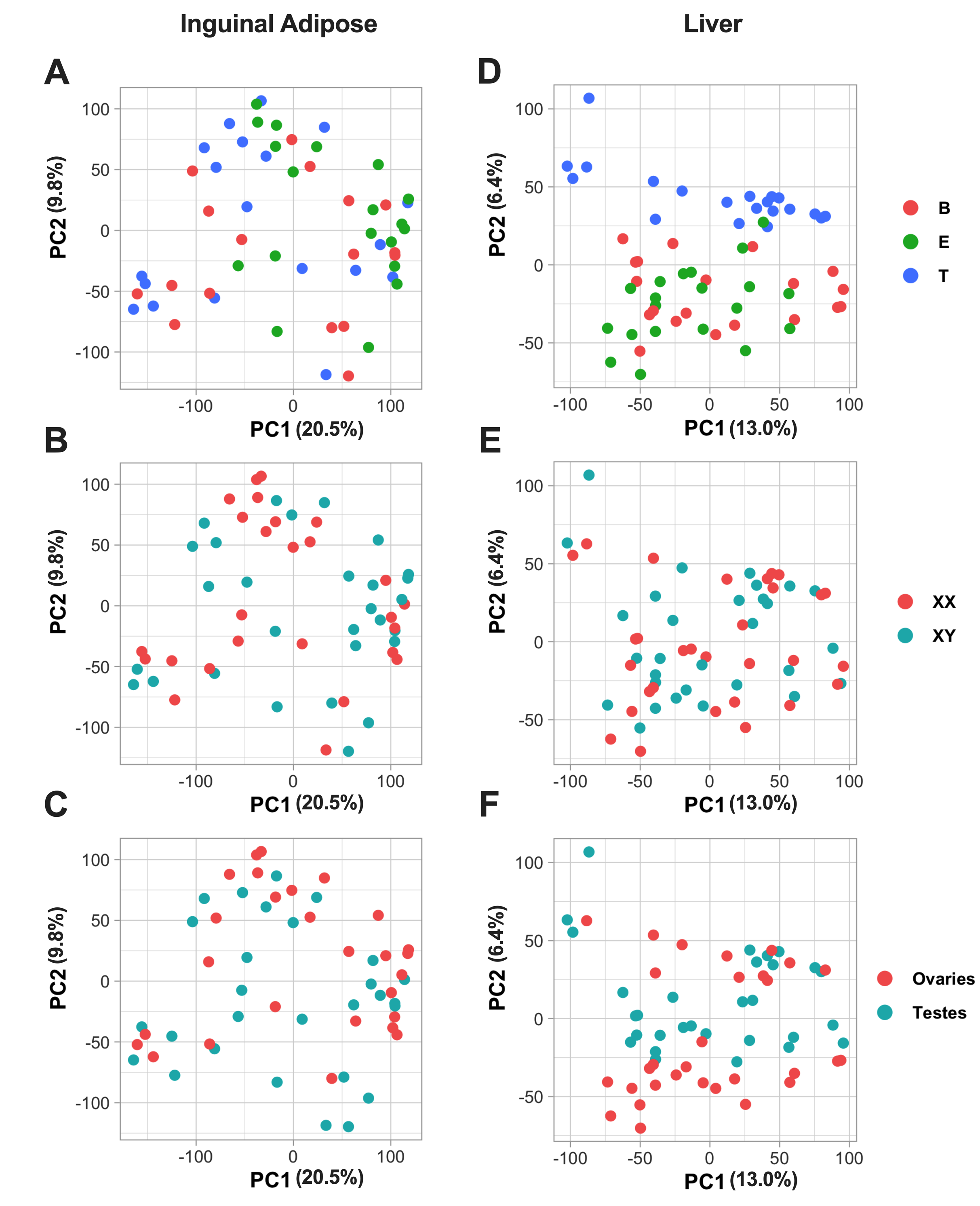

### Supplemental_Fig_S2.png

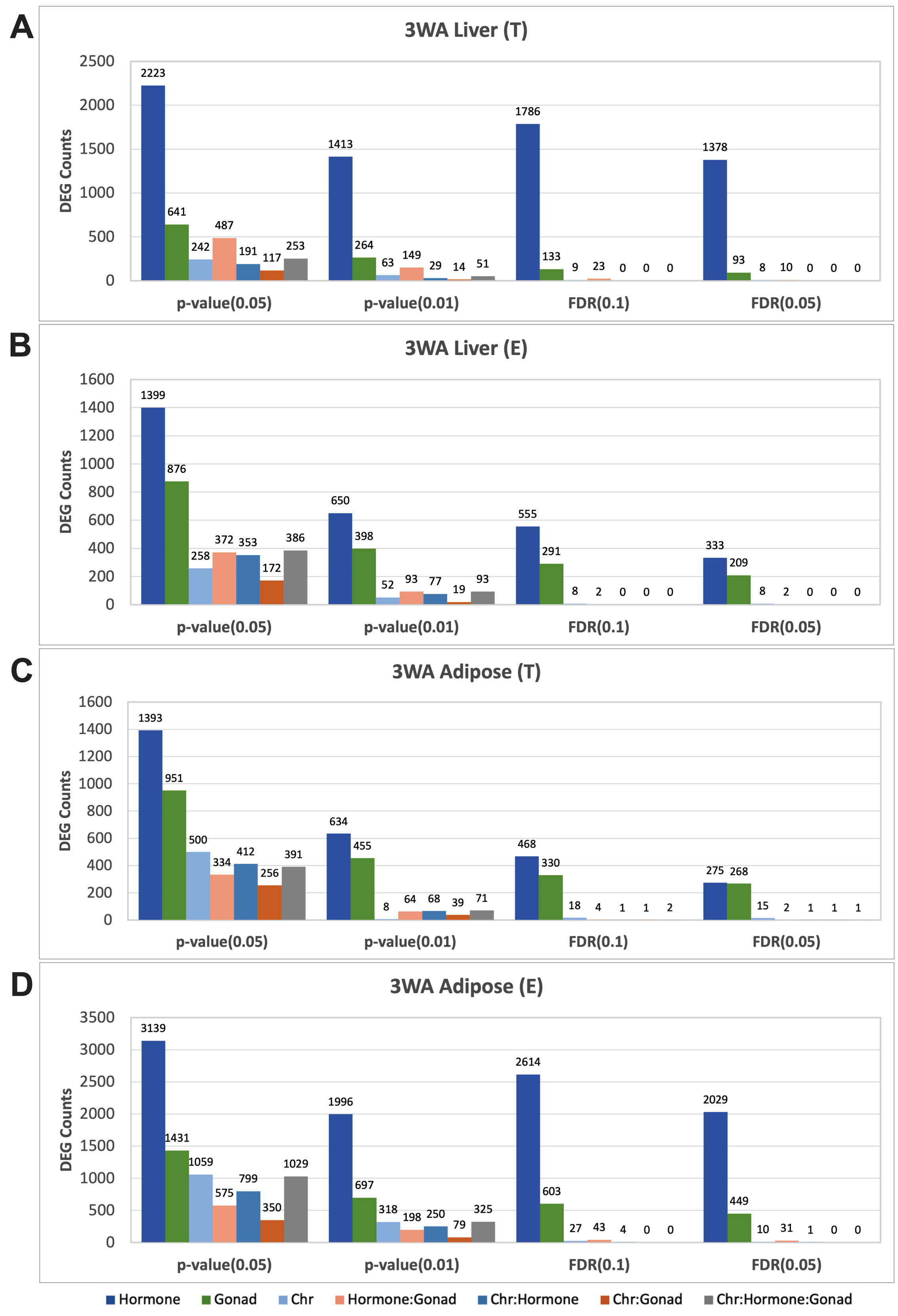

### Supplemental_Fig_S3.pdf

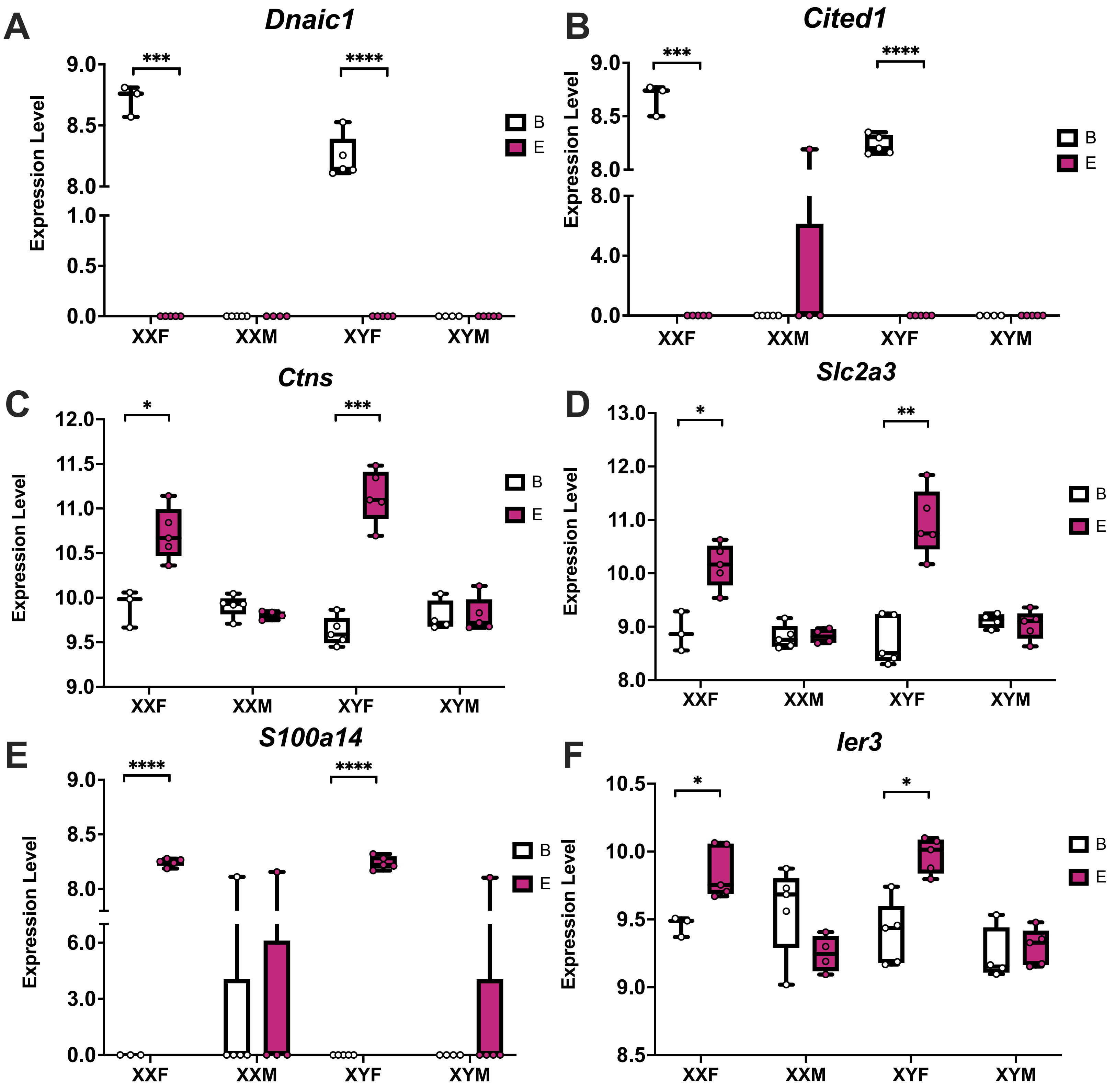

### Supplemental_Fig_S4.pdf

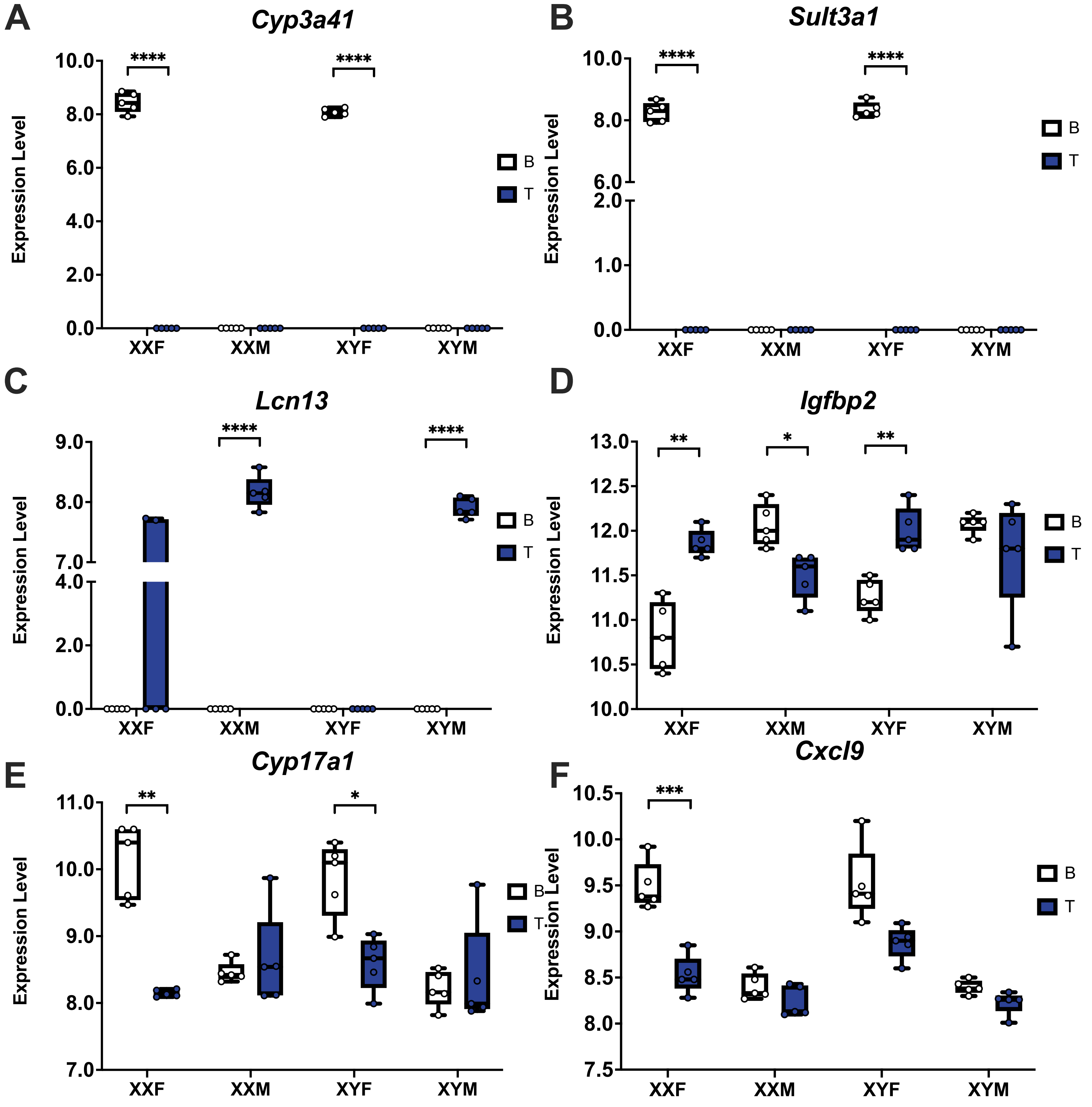

### Supplemental_Fig_S5.pdf

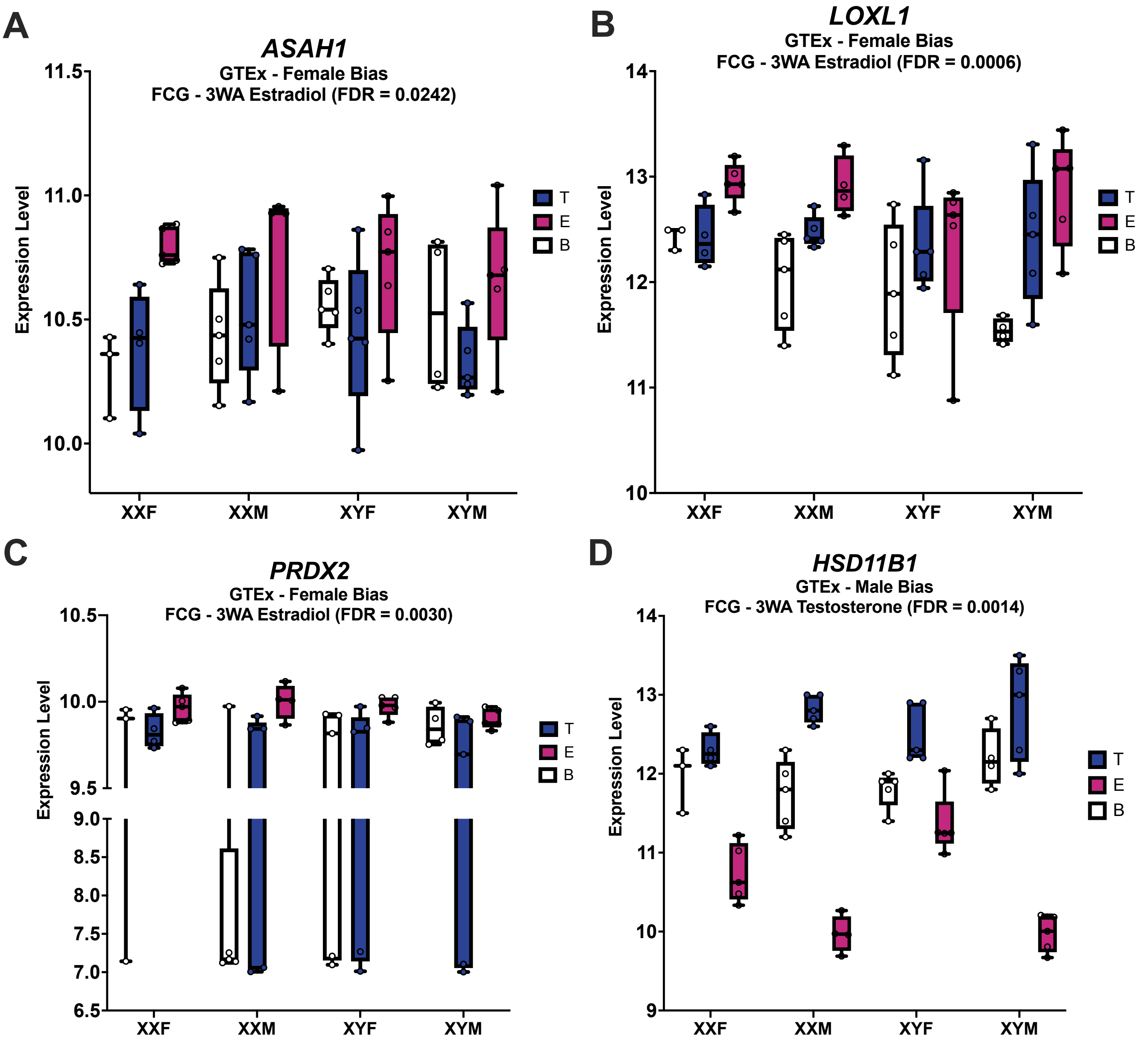

### Supplemental_Fig_S6.pdf

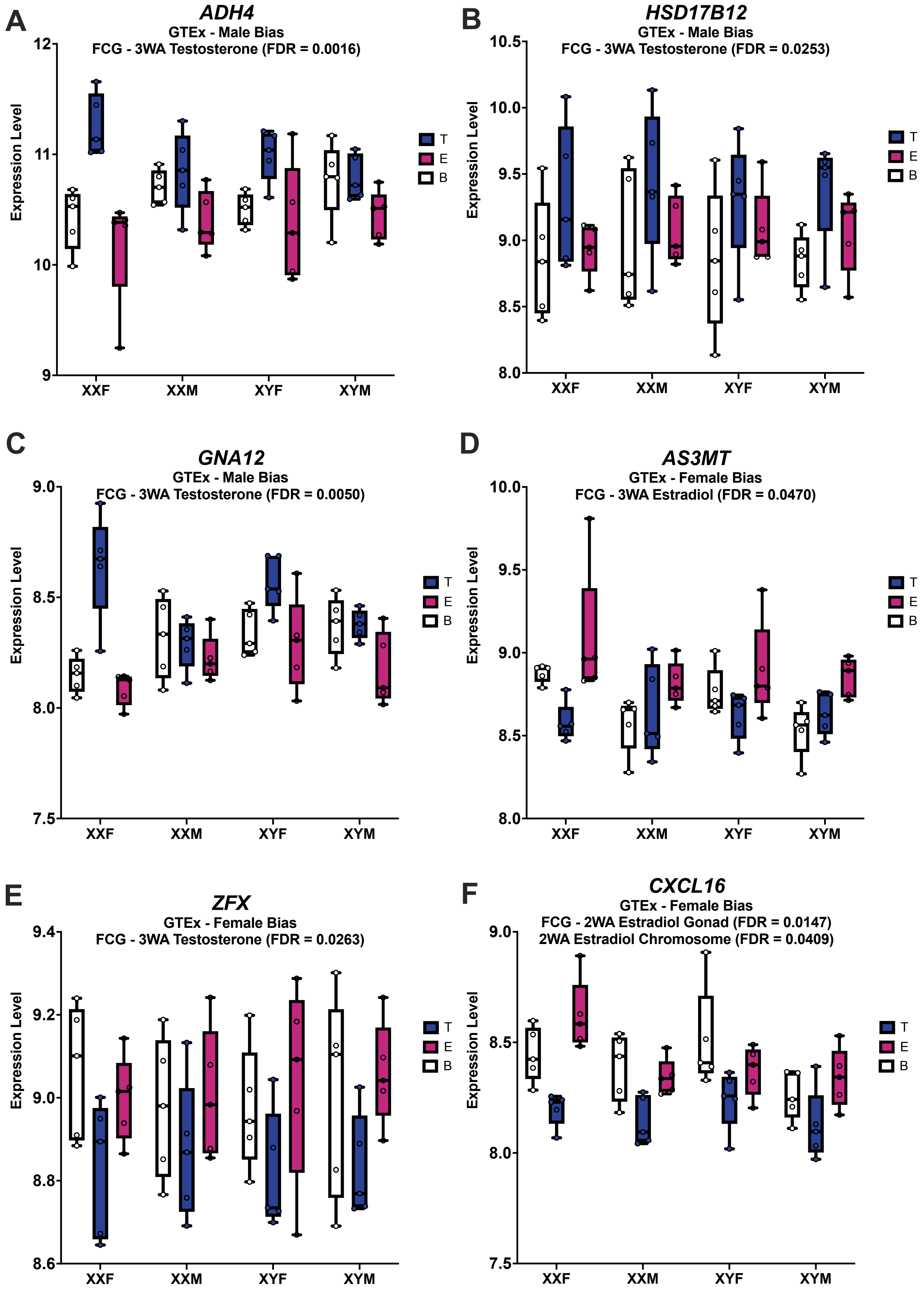

### Supplemental_Fig_S7.pdf

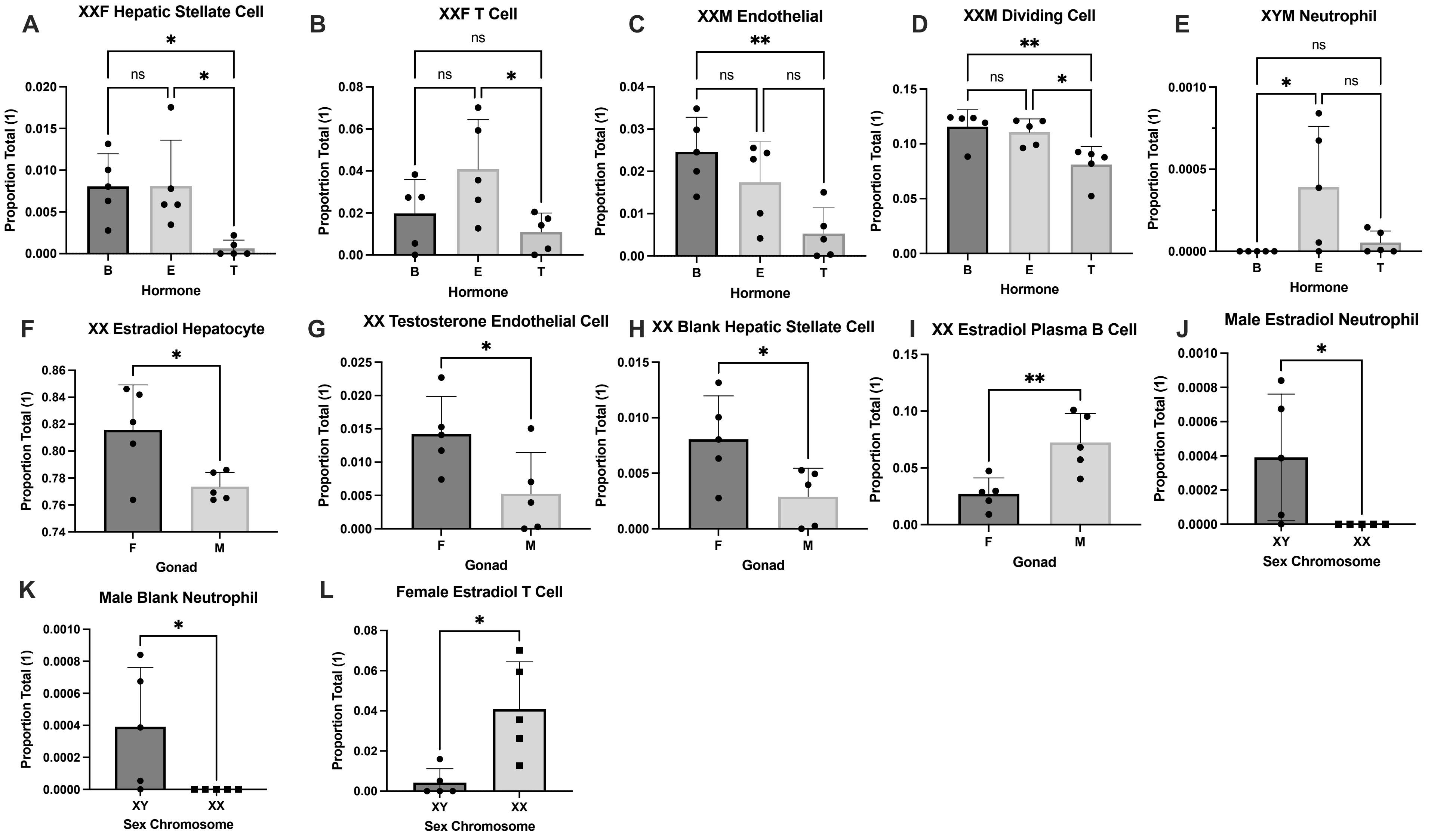

### Supplemental_Fig_S8.pdf

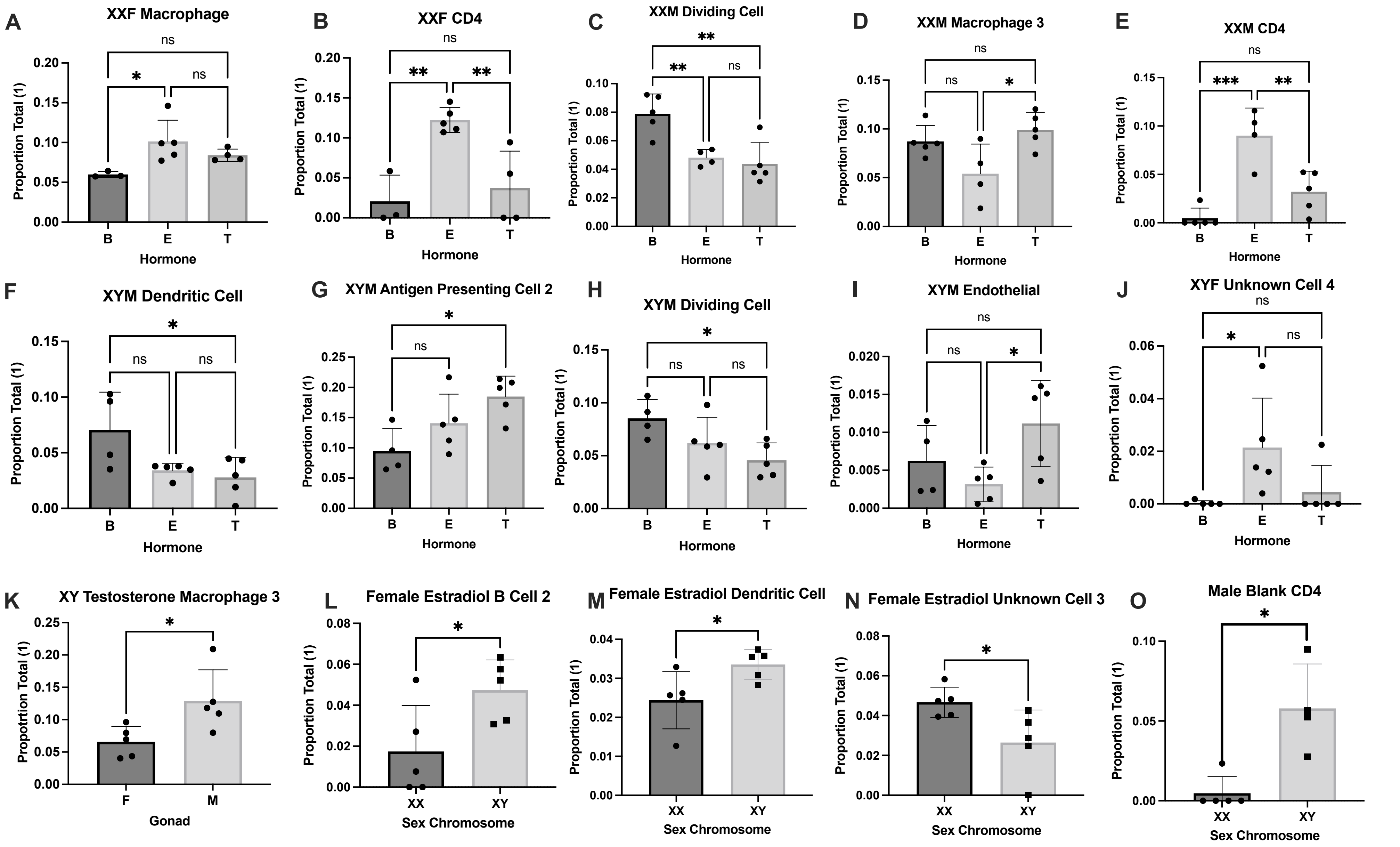
