## Supplementary material for "Relative contributions of sex hormones, sex chromosomes, and gonads to sex differences in tissue gene regulation": Supplemental Materials_Revision_Submission.docx

Arthur Arnold, Ph.D.

Department of Integrative Biology & Physiology, UCLA

610 Charles E Young Drive South

Los Angeles CA 90095-7239

Xia Yang, Ph.D.

Department of Integrative Biology & Physiology, UCLA

610 Charles E Young Drive South

Los Angeles CA 90095-7239

Supplemental Methods

***Overall Study Design***

Testosterone pellets contained 5mM testosterone (Steraloids, Inc., Newport, RI, USA) in a Silastic tube with 1.57mm interior diameter, 2.41mm outside diameter (Dow Corning, Midland, MI, USA) (Chen et al. 2013; Seney et al. 2013). These pellets produce testosterone levels in the high physiological range (Ngun et al. 2014). The estradiol pellets contained 50µg of 17beta-estradiol (Steraloids) dissolved in 25 µl sesame oil within a Silastic tube with interior diameter 1.98mm and 3.17mm outside diameter, sealed on the ends with 3mm of medical grade Silastic adhesive. The estradiol pellets provide sufficient estrogen to induce progestin receptors in the brain (Kudwa et al. 2009).

***Co-expression network construction and identification of co-expression modules affected by individual sex-biasing factors***

MEGENA splits gene clusters via divisive clustering based on the shortest path distance measure and a nested k-medoids clustering. At each step, the shortest path distance is minimized within each cluster, and the nested clustering process is repeated until no more compact child clusters can be identified. The Spearman correlation method with 100 permutations was implemented during the correlation analysis and p-value cut-off was set to 0.05. Module size was set from 50 to half of the vertices of planar filtered networks.

***Gene regulatory network analysis***

To predict potential regulators of the sex-biased DEGs, we used the Key Driver Analysis (KDA) function of the Mergeomics pipeline (Shu et al. 2016) and liver and adipose Bayesian networks (BNs). Firstly, BNs represent a class of gene regulatory networks, which can demonstrate the directed causal relationships between genes by using genetic and gene expression information of samples, as well as previously known regulatory relationships between genes. In the BNs, each edge is directed to a child node from a parent node, where the BN represents a multivariate probability distribution, and the state of each node is estimated by the states of its parent nodes. We leveraged the Reconstructing Integrative Molecular Bayesian Networks (RIMBANet) tool (Zhu et al. 2007; Zhu et al. 2008; Zhu et al. 2012) to construct tissue-specific BNs using both large human and mouse transcriptome and genome datasets for both adipose and liver (Yang et al. 2006; Wang et al. 2007; Emilsson et al. 2008; Schadt et al. 2008; Tu et al. 2012). Next, we used the KDA to predict key regulator, or key driver (KD), genes within the sex-bias DEG sets. The KDA uses a Chi-square like statistic, $\chi=\frac{O-E}{\sqrt{E}- \kappa}$ , to identify genes (KDs) that are connected to a significantly larger number of sex-bias genes than what is expected by random chance. O and E represent the observed and expected numbers of sex-bias genes in a hub subnetwork, and E is estimated by $\frac{N_{k}N_{p}}{N}$ where *N_p_* is the sex-bias gene set size, *N_k_* is the hub degree, and *N* is the full network order. The Benjamini-Hochberg method was used to correct *P*-values for the multiple hypothesis testing, and candidate genes with an FDR<5% were predicted as KDs of sex-bias DEGs. KDs were ranked based on their FDR scores and the top KDs were identified. Then, subnetworks of the top KDs were extracted in each tissue BN by collecting network-neighbors of the top KDs.

***Maker Set Enrichment Analysis (MSEA) to connect sex biasing DEGs with human diseases or traits***

To assess the potential role of the DEGs affected by each of the sex-biasing factors in human diseases, we collected the summary statistics of human GWAS for 73 diseases or traits that are publicly available via GWAS catalog (MacArthur et al. 2017). SNPs that have linkage disequilibrium of r^2^>0.5 were filtered to remove redundancies. To map GWAS SNPs to genes, we used GTEx Version 7 eQTL data for liver and adipose tissues (Lonsdale et al. 2013) to derive tissue-specific genes potentially regulated by the SNPs. We then use the MSEA function embedded in Mergeomics (Shu et al. 2016), which utilizes a modified chi-square statistic for the enrichment assessment in the sex-bias DEGs, which is summarized across a range of quantile-based cutoffs for the GWAS, instead of depending on a single GWAS *P*-value cutoff. This test analyzes the significance of enrichment for stronger disease-GWAS *P*-values, by comparing the GWAS *P*-values of a given eSNP set against the eSNP sets that were mapped from randomly generated gene sets. Our MSEA approach is based on a set of quantile-based cutoffs and is not based on a single GWAS *P*-value cutoff; hence it avoids artefacts while producing more stable enrichment scores. The definition of the modified chi-square statistics used in the MSEA is:
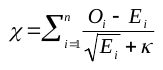
$,$ where *O* and *E* are the numbers of the observed and estimated positive findings, respectively (i.e. findings above the *i*-th quantile point); n is the number of quantile points (10 points were identified ranging from the top 50% to top 99.9% signals based on the GWAS *P*-value rankings),. For the MSEA procedure, an FDR<5% cut-off was used, which is estimated by the Benjamini-Hochberg method (Benjamini and Hochberg 1995) to identify significantly enriched sex-bias DEGs for the disease-GWASs.

**Supplemental Figures**

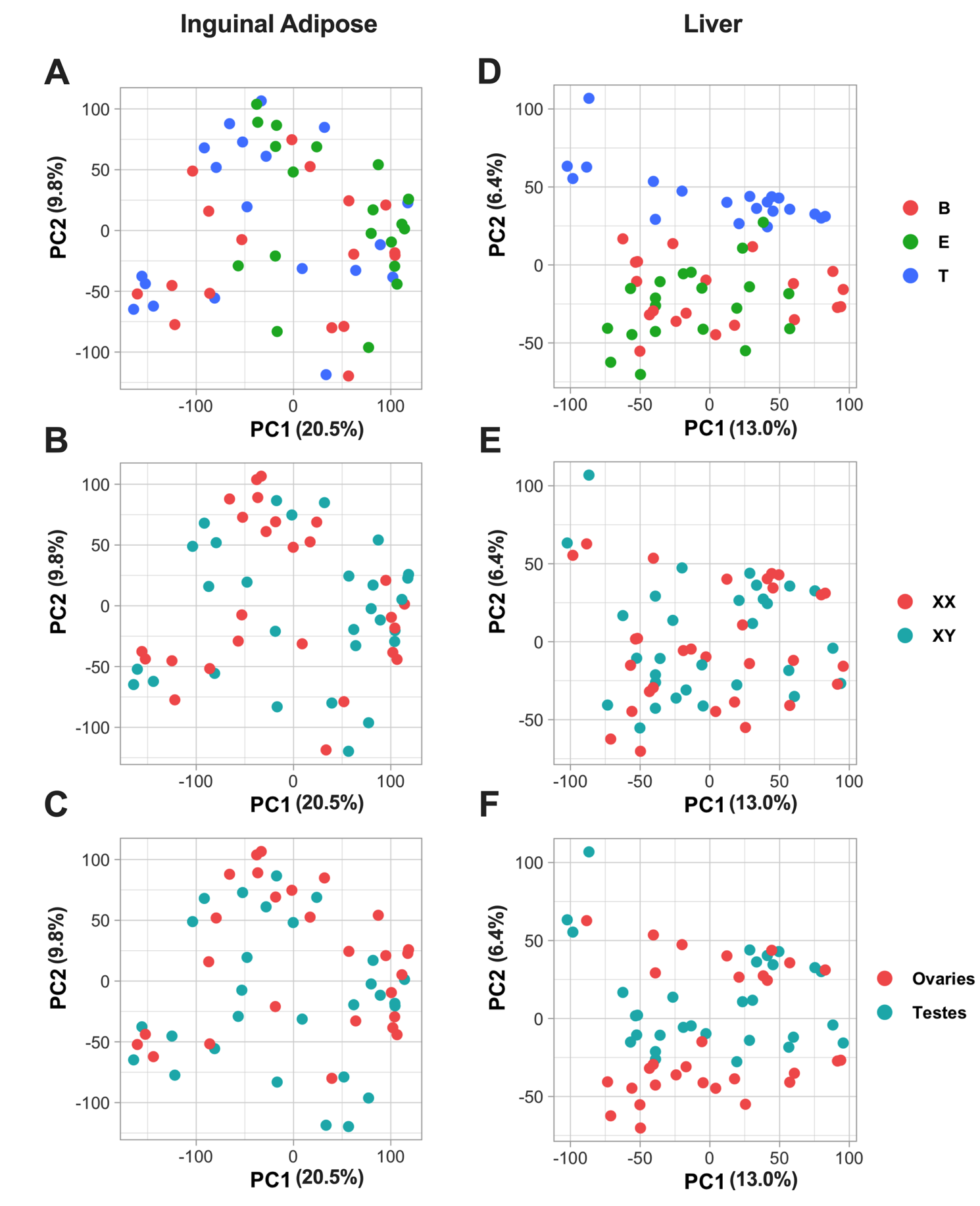

Supplemental Figure S1**: PCA plots of inguinal adipose and liver samples colored by different factors**

**A.** Inguinal adipose tissue PCA plot, with samples labelled by hormone types - blank (red), estradiol (green), and testosterone (blue). **B.** Inguinal adipose tissue PCA plot, with samples were labelled by sex chromosome categories - XX (red) and XY (blue). **C**. Inguinal adipose tissue PCA plot, with samples labelled by gonadal sex categories - ovaries (red) and testes (blue). **D.** Liver tissue PCA plot, with samples were labelled by hormone types - blank (red), estradiol (green), and testosterone (blue). **E**. Liver tissue PCA plot, with samples labelled by sex chromosome categories - XX (red) and XY (blue). **F**. Liver tissue PCA plot, with samples labelled by gonadal sex categories - ovaries (red) and testes (blue).

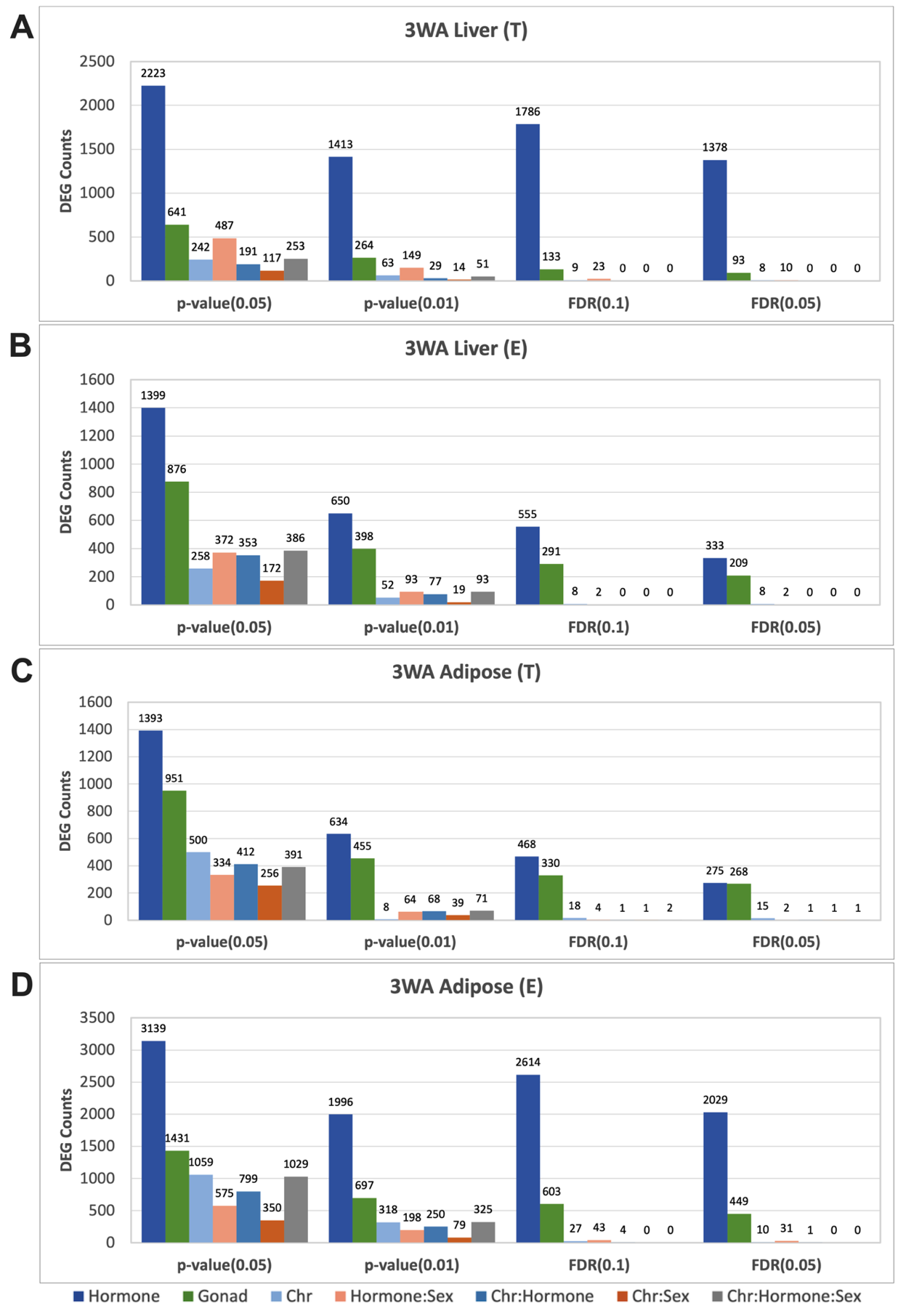

Supplemental Figure S2**:** Bar graphs representing the 3-way ANOVA DEG numbers across various statistical cutoffs. Across the cutoffs there is a consistent trend of which sex biasing factor produces greater influence on DEG numbers: hormones (testosterone T or estradiol E) > gonad > sex chromosome. (A) Liver 3WA DEG counts when Testosterone (T) and blank groups are considered. (B). Liver 3WA DEG counts when Estradiol (E) and blank groups are considered. (C) Adipose 3WA DEG counts when Testosterone (T) and blank groups are considered. (D) Adipose 3WA DEG counts when Estradiol (E) and blank groups are considered.

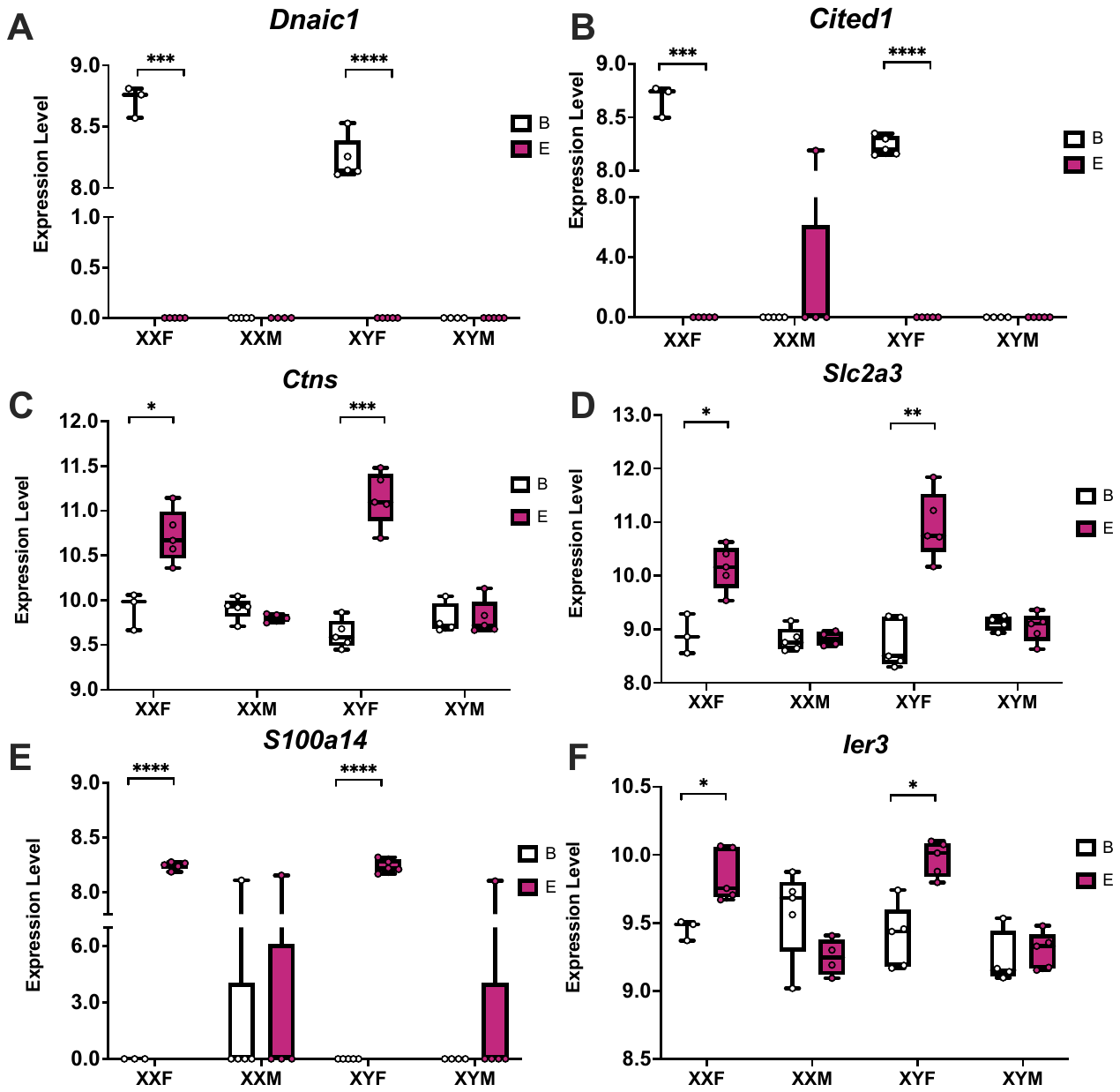

Supplemental Figure S3**:** Bar Plots highlighting genes that showcase significant interactions between estradiol and gonad type in adipose tissue. T-test was used to calculate within genotype statistical significance. FDR<0.0001****, FDR<0.001***, FDR<0.01**, FDR<0.05*

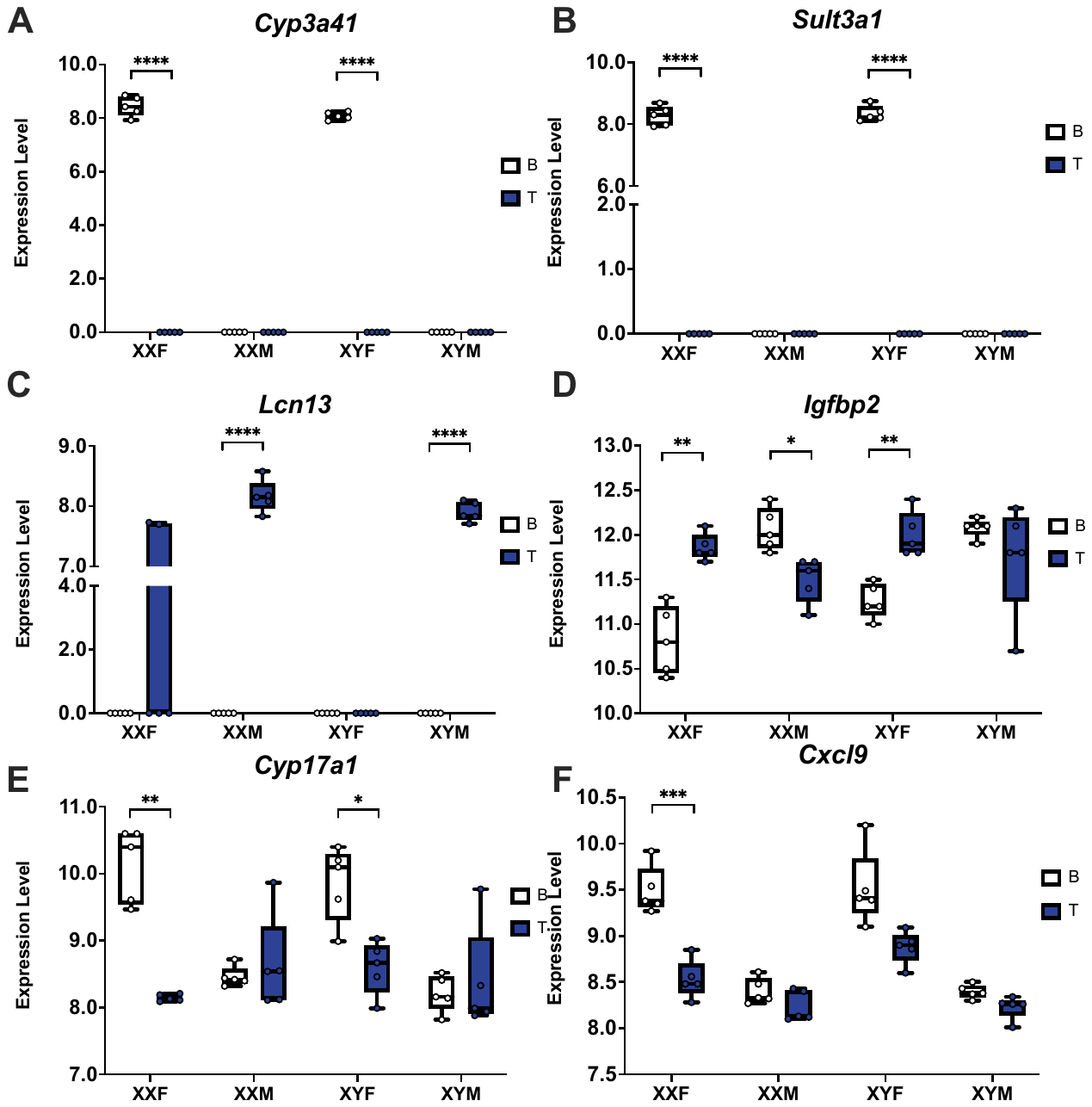

Supplemental Figure S4**:** Bar Plots highlighting genes that showcase significant interactions between testosterone and genotypes in liver. T-test was used to calculate within genotype statistical significance. FDR<0.0001****, FDR<0.001***, FDR<0.01**, FDR<0.05*

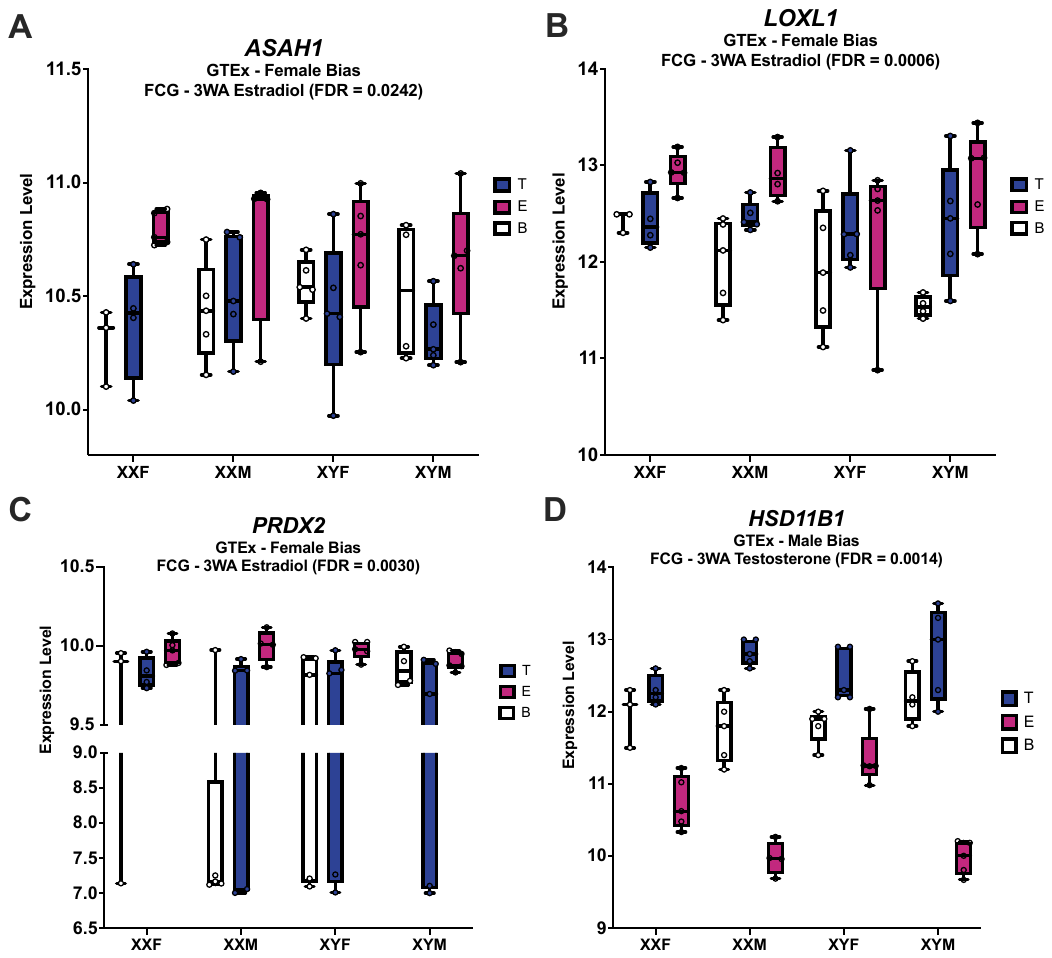

Supplemental Figure S5**:** Bar Plots highlighting genes that showcase sex bias in the human GTEx study (Oliva et al. 2020) which also show a matched sex bias through one of the sex biasing factors in adipose tissue. We highlight in the heading of each panel the statistical test conducted and their associated FDR value.

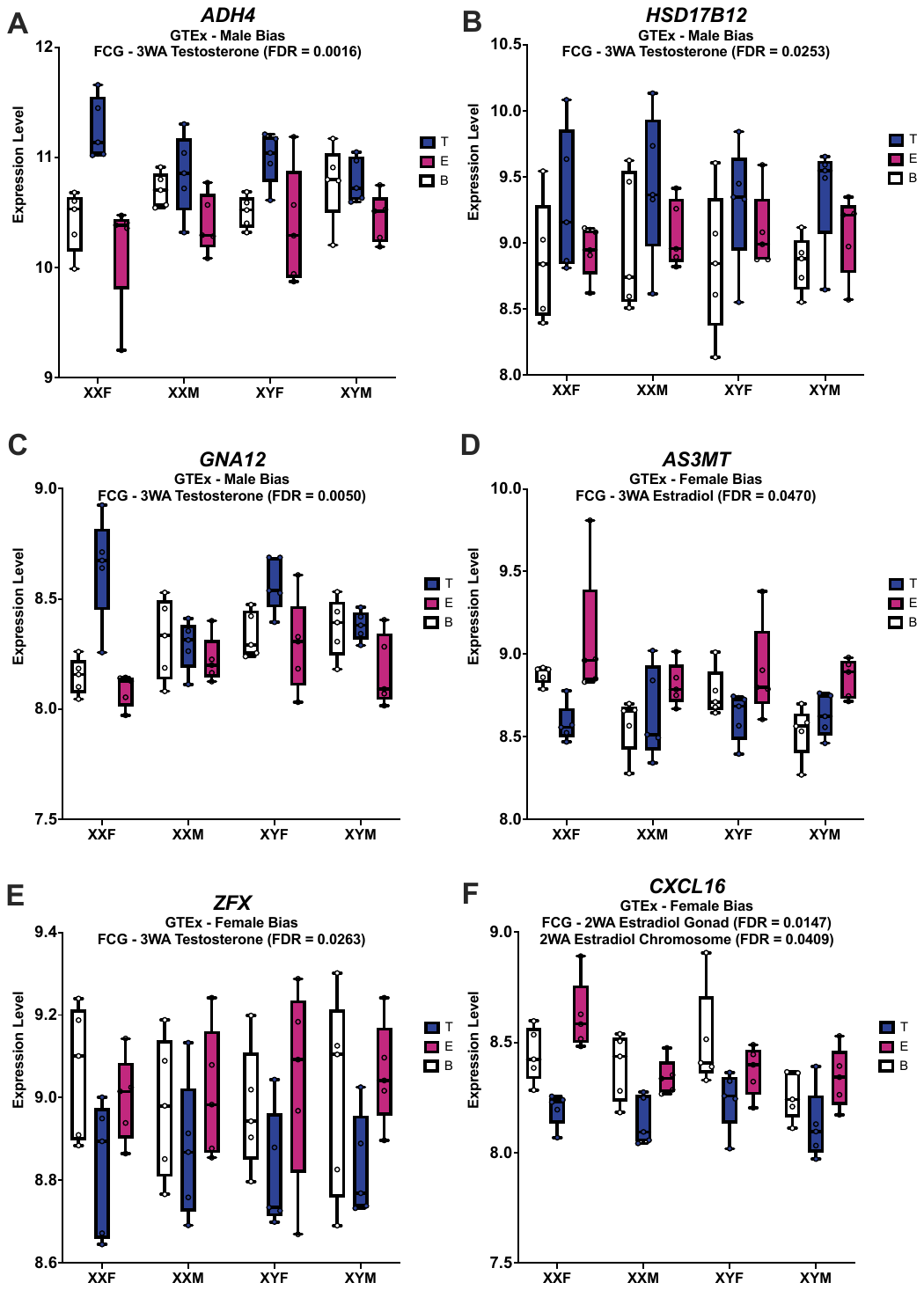

Supplemental Figure S6**:** Bar Plots highlighting genes that showcase sex bias in the human GTEx study (Oliva et al. 2020) which also show a matched sex bias through one of the sex biasing factors in liver tissue. We highlight in the heading of each panel the statistical test conducted and their associated FDR value.

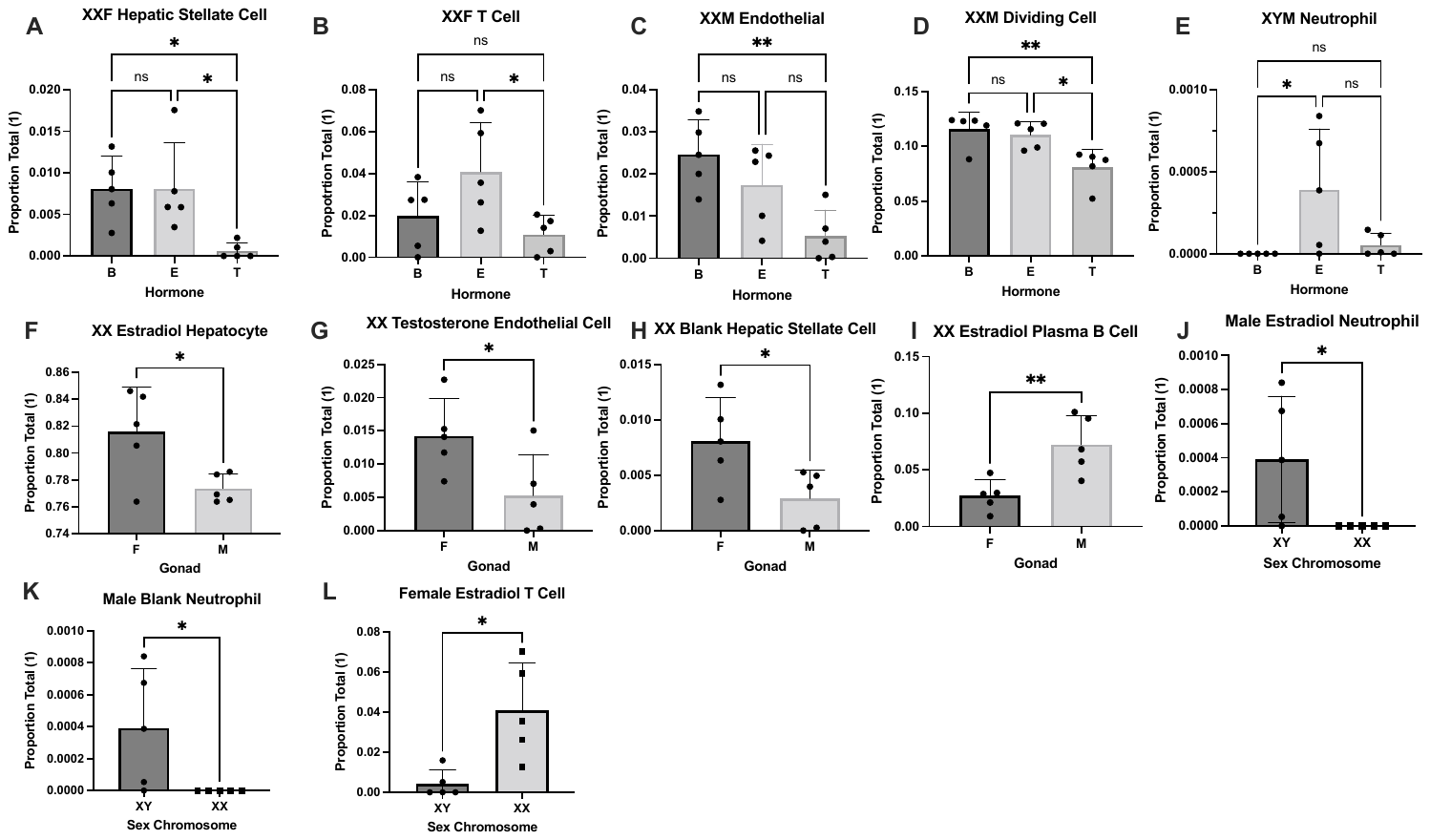

Supplemental Figure S7**:** Deconvolution results for the liver, highlighting cell types that showed a statistical difference in cell type proportion between hormone treatments (A-E), gonads (F-I), and sex chromosome (J-K). * FDR<0.05, ** FDR <0.01, ns. not significant. For hormonal comparison we used the 1 Way ANOVA followed by post hoc analysis and for gonad and chromosome we used a T-test to calculate statistics.

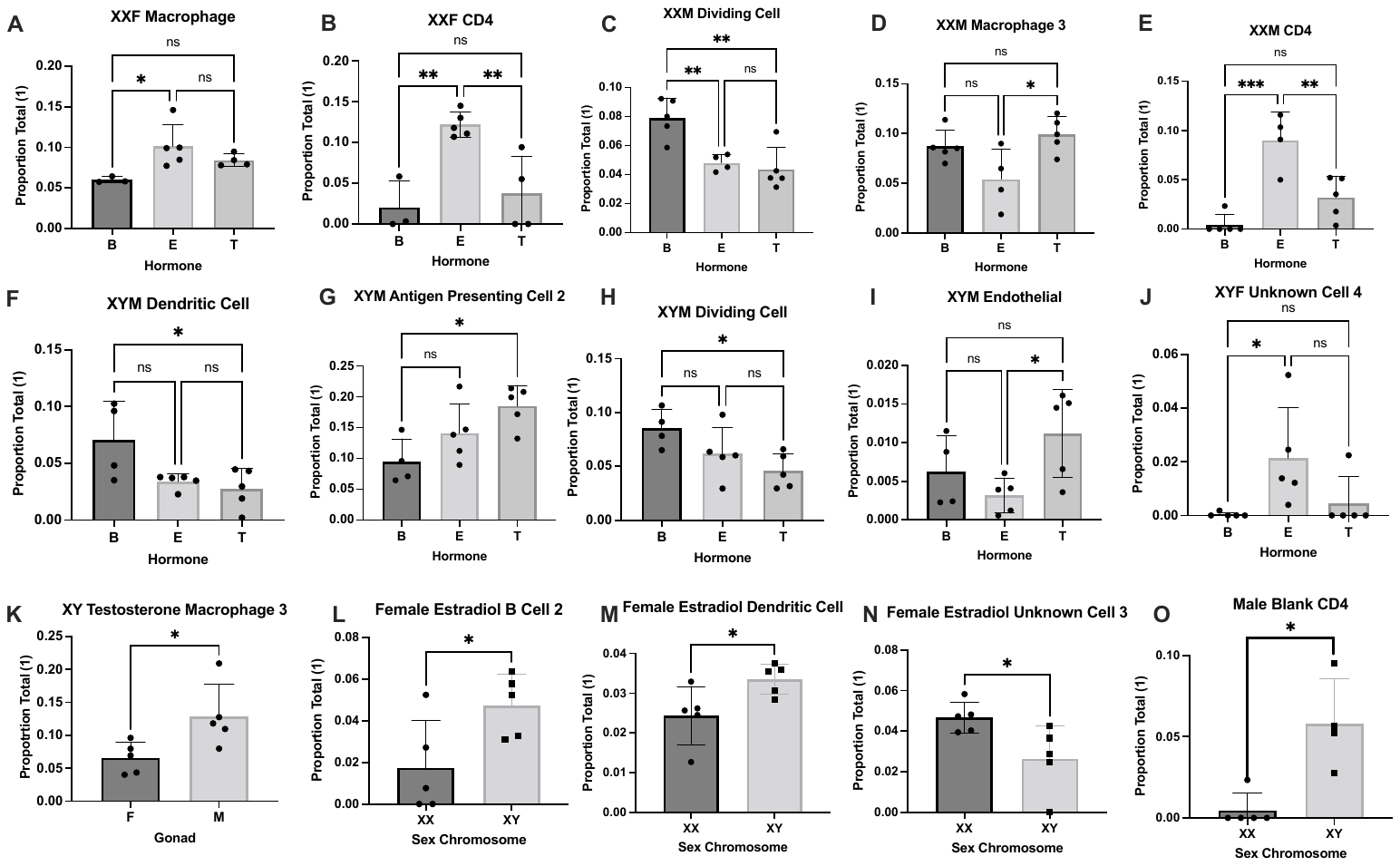

Supplemental Figure S8**:** Deconvolution results for the adipose tissue, highlighting cell types that showed a statistical difference in cell type proportion between hormone treatments (A-J), gonads (K), and sex chromosome (L-O). * FDR<0.05, ** FDR <0.01, ns. not significant. 1 Way ANOVA followed by post hoc analysis to calculate statistics for those with three groups and T-test was used for those with two groups.

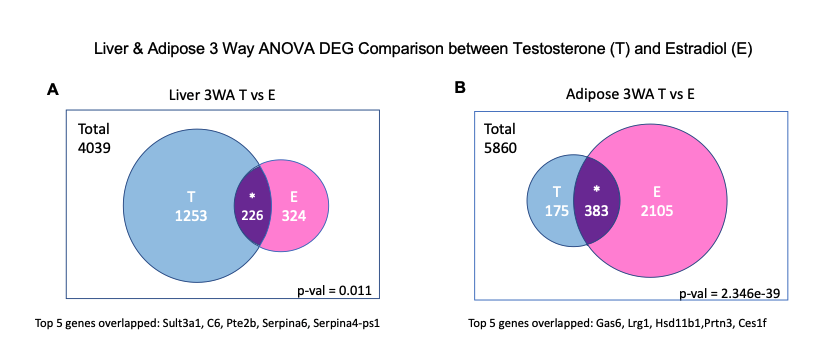

Supplemental Figure S9**:** Venn diagrams representing 3-way ANOVA DEG comparison between testosterone (T)- and estradiol (E)-treated group in liver and inguinal adipose tissue.

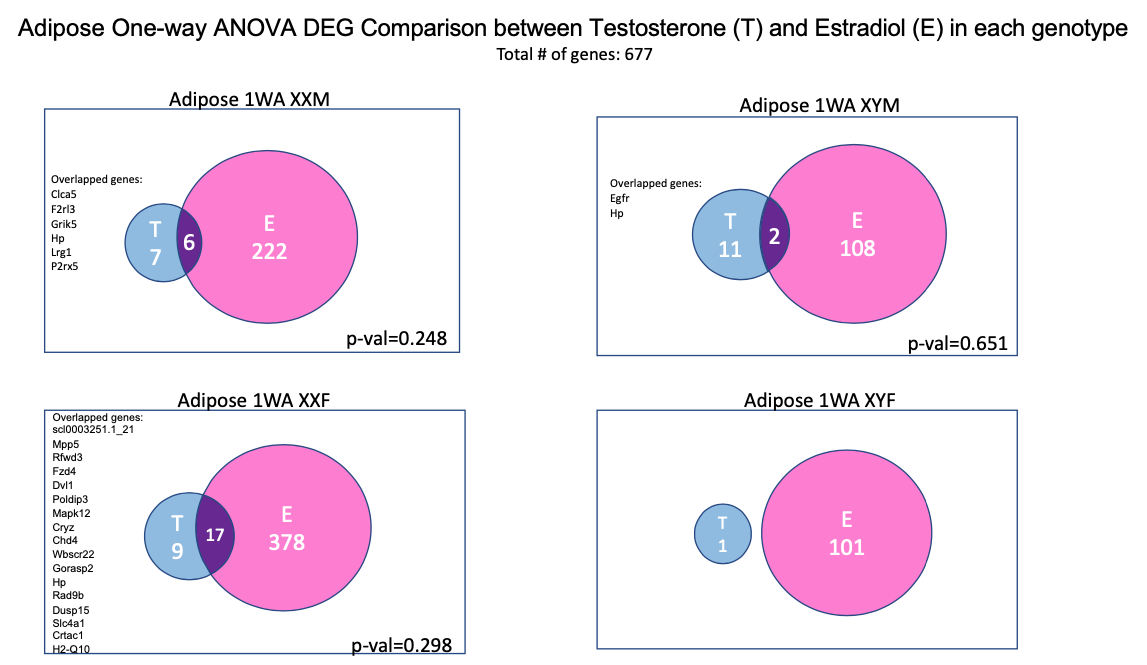

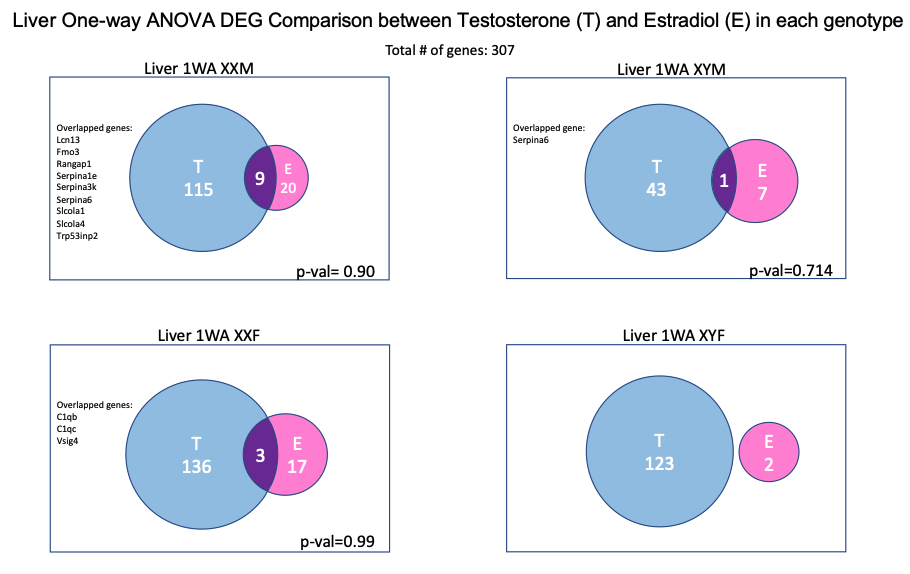

Supplemental Figure S10**:** Venn diagrams representing 1-way ANOVA DEG comparison between testosterone (T)- and estradiol (E)-treated group across all genotypes in liver and inguinal adipose tissue.

Supplemental Tables

Tables S1-S19 are available as separate online Excel files.
